# Rhizaria are unexpectedly abundant and exhibit taxonomic and trophic diversity in the eastern subarctic Pacific

**DOI:** 10.1101/2025.05.06.652060

**Authors:** Jaime R. Blais, Suzanne L. Strom

**Affiliations:** Shannon Point Marine Center, Western Washington University, Anacortes, WA 98221

## Abstract

Rhizaria are a diverse supergroup of large marine protists that are often overlooked due to their fragility, lower abundances, and wide size range relative to other plankton. Despite their global distribution, Rhizaria ecology and biogeography is poorly understood due to a paucity of datasets and use of differing methodologies. Here we present the first characterization of Rhizaria ecology in the northern Gulf of Alaska (NGA), a variable yet productive subarctic ecosystem with important fisheries that is experiencing long-term warming. Seawater samples were collected from CTD-secured Niskin bottles at stations within the NGA Long-Term Ecological Research study area during summer 2023. We report some of the highest Rhizaria abundances (25 cells L^-1^) from any ocean environment to date and thus suggest a restructuring of the current biogeographical paradigm that posits highest abundances at the equator and decreases at higher latitudes. Acantharia was the most ubiquitous subgroup. Distinct depth niches were also revealed: Foraminifera dominated surface waters, Radiolaria exhibited a cosmopolitan distribution, and Phaeodaria were the deepest living. Prey captures and algal interactions primarily occurred offshore in the upper water column. A wide range of taxa had captured prey while the hosts to presumptively symbiotic algae were mainly Foraminifera and Acantharia. We highlight Rhizaria as key players in NGA food web dynamics as evidenced by their wide depth distributions, taxonomic diversity, and variable nutrition strategies.

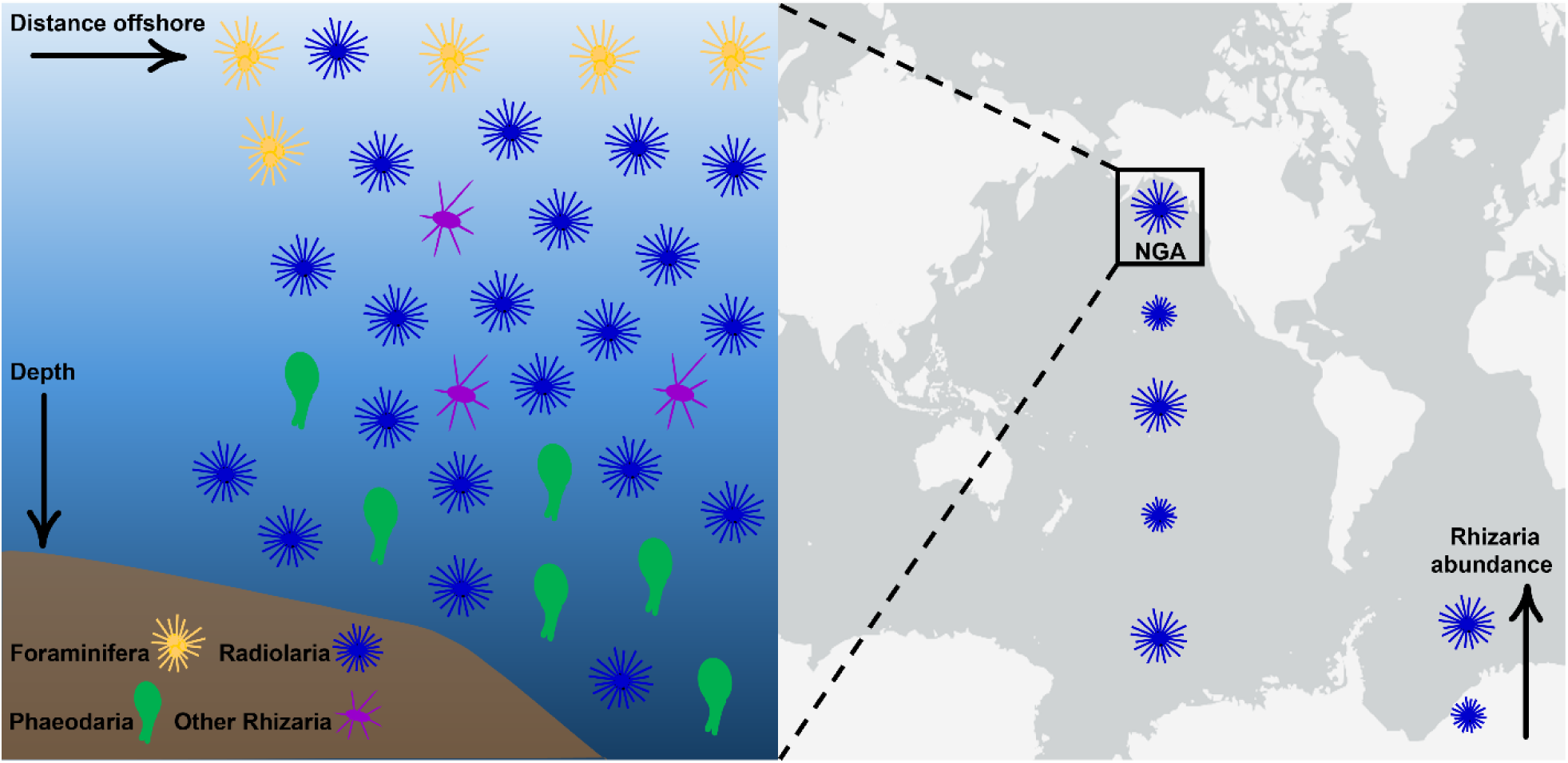
Graphical Abstract. Distribution of Rhizaria subgroups in the northern Gulf of Alaska (left). Proposed revision of the biogeographical distribution of Rhizaria in the Pacific and Southern Oceans (right).

**Highlights:**

- The N. Gulf of Alaska contains some of the highest Rhizaria abundances yet reported
- Acantharia was the most abundant taxon
- Rhizaria subgroups inhabited distinct depth niches
- A wide range of taxa had captured prey
- Foraminifera and Acantharia were the most common hosts to algal cells

## INTRODUCTION

A supergroup of large (roughly 50 µm to 5 mm) amoeboid hetero- and mixotrophic marine protists, Rhizaria’s unique features include pseudopodal cytoplasmic projections for feeding and intricate skeletal structures made of silica (Polycystina and Phaeodaria), calcium carbonate (Foraminifera), or strontium sulfate (Acantharia). Although Rhizaria are found globally in marine ecosystems, the fundamentals of their biology and ecology are poorly understood. We sought to investigate their importance in the northern Gulf of Alaska Long-Term Ecological Research area (NGA-LTER). This productive subarctic region extends about 200 km off the south-central coast of Alaska (Figure 1), spanning the continental shelf to the open ocean. The highly seasonal ecosystem supports a semi-predictable spring phytoplankton bloom, considerable copepod and gelatinous zooplankton biomass, and multiple commercially important fish populations like walleye pollock (Dorn et al. 2017, Strom 2023). Only a few Rhizaria studies have taken place in the Gulf of Alaska, all at station PAPA considerably further south (50°N 145°W). These were exclusively vertical flux analyses (Takahashi 1987, 1997) that did not address the ecology of living planktic Rhizaria.

**Figure 1.**
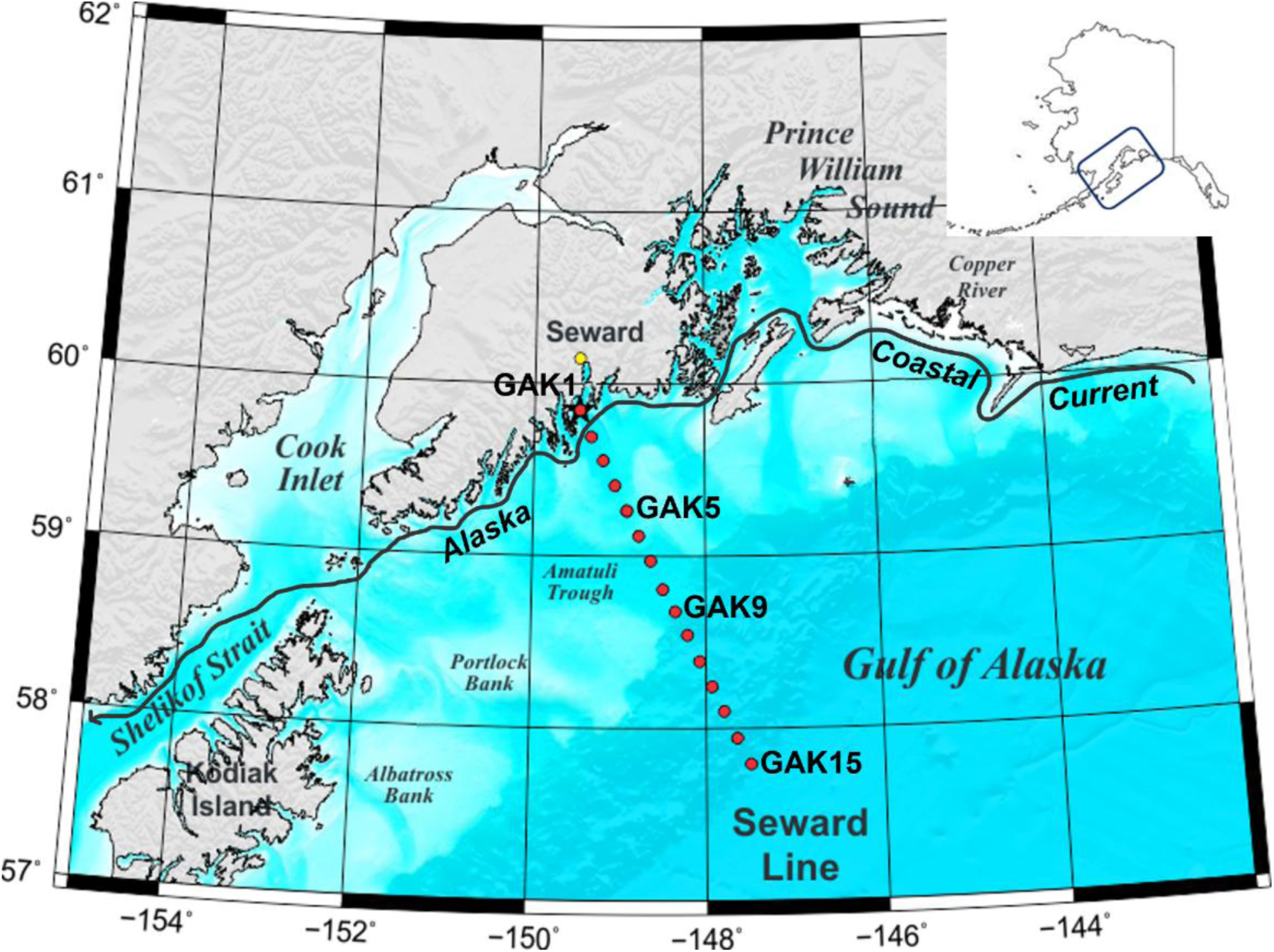
Northern Gulf of Alaska Long-Term Ecological Research study region. Samples were collected at four stations along the Seward Line: GAK1, GAK5, GAK9, and GAK15.

Rhizaria are now recognized as important players in the ocean’s biological carbon pump, biogeochemical cycling, and trophodynamics. Due to their large sizes, inorganic skeletons, and global distribution these amoeboid protists, especially the large and colonial species, facilitate particle ballasting and carbon and biogenic silica export (Lampitt et al. 2009, Guidi et al. 2016, Biard et al. 2018, Stukel et al. 2018, Gutierrez-Rodriguez et al. 2019, Ikenoue et al. 2019). They also connect microbial and protist networks to higher trophic levels as predators of phytoplankton and microzooplankton, prey for macrozooplankton and fish, and host to algal cells (Biard 2022). Using sticky pseudopodia, Rhizaria feed opportunistically on a variety of planktonic organisms (Swanberg and Caron 1991). There have only been a handful of reports on Rhizaria as prey themselves, mainly from gut content analysis. Known predators include crustaceans such as mysids, euphausiids, amphipods, and copepods (Hopkins 1985, 1987, Gowing and Wishner 1986), salps (Hopkins and Torres 1989, Gowing 1989), liparid fish (Takami and Fukui 2012) and deep-sea smelt (Hopkins and Torres 1989).

In addition to feeding, many surface-dwelling Radiolaria and Foraminifera host symbiotic algae such as cyanobacteria (*Synechococcus* sp. and *Prochlorococcus* sp.), haptophytes, and dinoflagellates, making them mixotrophs (Decelle et al. 2015). Photosymbionts provide Rhizaria with carbon for use in maintenance metabolism or growth (Swanberg 1983, Stoecker et al. 2017). Mixotrophic protists help sustain food webs and may aid in resilience in resource-limited regions because they are capable of transitioning between nutrition strategies depending on environmental conditions or prey availability (Stoecker et al. 2017, Strom et al. 2024). The offshore NGA waters are typically high in nitrate but are iron-limited, resulting in low phytoplankton biomass or chlorophyll. These HNLC conditions restrict primary productivity and availability of phytoplankton prey, especially larger cells (Nishioka et al. 2021). Therefore, Rhizaria-algal relationships likely support the NGA food web particularly in offshore waters.

Biogeographical and bathymetric faunal zones or provinces have been assigned to Radiolaria, Foraminifera, and Phaeodaria species many times in an attempt to categorize the distributions of these relatively rare protists; important influences on their biogeographical distribution appear to be latitude, temperature, and/or water mass movements (Casey 1966, 1971, Kling 1966, Renz 1976, Bé 1977, Boltovskoy and Correa 2016, 2017) as well as nutrient concentrations and primary productivity (Boltovskoy et al. 2017). However, despite the ubiquity of Rhizaria, our understanding of their ecology and biogeographical patterns is poor for multiple reasons. Abundances are low compared to more widely studied plankton and sampling challenges exist due to the group’s wide size range and biomineralized skeleton fragility (Suzuki and Not 2015, Boltovskoy et al. 2017). Since the time of the 1870s Challenger Expedition, when the first seminal works on Rhizaria ecology and diversity were published, most sea-going scientists have utilized net tow or sediment trap collection methods to measure water column distributions and flux (e.g. Haeckel 1887, Kling 1979, Morley and Stepien 1985, Boltovskoy and Riedel 1987, Takahashi 1987, Bernstein et al. 1990, Okazaki et al. 2005, Ishitani and Takahashi 2007, Ikenoue et al. 2012). The former can disrupt fragile skeletons and misses cells smaller than the mesh size (Michaels 1988, Stoecker 1996), while the latter captures only forms that sink intact. More recently, *in situ* imaging instruments like the Underwater Vision Profiler, FlowCAM, and Zooscan have been used to estimate abundances, biomass, and flux of large Rhizarians (>600 µm, Biard et al. 2016, 2018, Stukel et al. 2018, Biard and Ohman 2020; >200 µm Llopis Monferrer et al. 2022), omitting cells in the lower portion of the size range. Our goal was to capture the entire size range and not discriminate against sensitive taxa. Therefore, we employed a meticulous protocol involving Niskin bottle water collection, reverse concentration by gentle siphoning, and formalin-strontium fixation, an approach inspired by Gowing and Garrison (1992) and Stoecker et al. (1996).

This study addresses a large gap in our understanding of Rhizaria ecology in the North Pacific Ocean and, more broadly, Acantharia and Taxopodida ecology worldwide. The majority of Rhizaria plankton research to date has focused on Polycystine Radiolaria, Phaeodaria, and Foraminifera, while global data on Acantharia and Taxopodida are scarce (e.g. Casey 1966, Bé and Tolderlund 1971, Kling 1979, Kling and Boltovskoy 1995, Welling 1992, Takahashi 1997, Okazaki et al. 2004, Itaki et al. 2008, Ikenoue et al. 2019). Taxopodida have been overlooked to an even greater extent than Acantharia; this may be because for many years their membership in the Rhizaria supergroup was contested, and they were instead classified as Heliozoans (Cachon and Cachon 1978). Now considered to be Radiolarians on the basis of phylogenetic and evolutionary analyses (Cavalier-Smith 1993, Nikolaev et al. 2004, Krabberød et al. 2011, 2017), Taxopodida have been found worldwide where they have been explicitly included in sample analysis (e.g. Southern Ocean, Gowing 1989, González 1992, Gowing and Garrison 1992; equatorial Pacific, Takahashi and Ling 1980; Eastern Indian Ocean, Munir et al. 2020, 2021; and Norwegian fjords, Ikenoue et al. 2023).

Given their demonstrated importance to the oceanic food web and carbon pump in other systems, we sought to better understand the role of Rhizaria in a previously unstudied subarctic region of the North Pacific, the NGA, through investigation of their ecology, biology, and diversity. We present a comprehensive summary of Rhizaria ecology that includes the first quantitative analysis of summer abundances and biomass in the context of cross-shelf and depth distributions, along with community composition and depth-niche analyses, morphotypes, light microscope imagery, and incidences of prey capture and algal cell interaction. In addition to addressing prevalence, diversity, and trophic mode in the northern reaches of the North Pacific, our study addresses global gaps in knowledge of under-sampled Rhizarians such as Acantharia and Taxopodida. Finally, this detailed addition to the worldwide Rhizarian database allowed us to propose revisions to current paradigms concerning global Rhizaria biogeography.

## METHODS

### Field sampling

This research project was conducted as part of the northern Gulf of Alaska Long-Term Ecological Research Program (NGA LTER). Seawater samples were collected between June 29 and July 6, 2023 on R/V *Kilo Moana* from CTD-secured Niskin bottles along the Seward transect line at “Gulf of Alaska” stations GAK1, GAK5, GAK9, and GAK15 (Figure 1). Four 35.25 L “depth interval” samples were collected at each station. Each sample consisted of water from three combined 12 L Niskin bottles; each Niskin contained water from a different depth for a total of three combined depths per depth interval sample (e.g. the 0-20 m sample comprised water from 0, 10, and 20 m; see Supplemental Table 1). Chosen depth intervals depended on bottom depth at station but together represented most of the water column. Seawater was gently transferred from Niskin bottles into buckets with lids, then reverse concentrated with siphoning through a 50 µm mesh sieve at 10°C, as modeled after Gowing (1989) and Stoecker et al. (1996). Each ∼35 L sample was concentrated in two stages to a final volume of 400 mL, fixed with 2% formalin (20% formaldehyde buffered with 100 g L^-1^ hexamethylenetetramine) + 0.16 mg mL^-1^ strontium chloride (Stoecker et al. 1989, 1996), and stored at 4°C. This concentration of SrCl_2_ (10x that of normal seawater) was used to prevent dissolution of Acantharian skeletons (Beers and Stewart 1970). The Niskin bottle collection method was chosen as the best approach to quantitatively sample Rhizaria on the basis that plankton net tows can underestimate abundances and damage fragile skeletons, especially those of Acantharians (Michaels 1988, Gowing 1989, Michaels et al. 1995, Stoecker et al. 1996).

Oceanographic parameters were measured at every GAK station along the Seward Line. Salinity and temperature were measured with Sea-Bird SBE 4C conductivity and SBE 3P temperature sensors, respectively. Size-fractionated chlorophyll-a concentrations (<20 µm, >20 µm) were measured at 0, 10, 20, 30, 40, 50, and 75 m as described in Strom et al. (2016). Beam transmission was measured with a Wetlabs C-Star 25 cm transmissometer. Nitrate concentrations were measured at the University of Alaska Fairbanks Nutrient Analytical Facility following procedures outlined in the GO-SHIP Repeat Hydrography Nutrient Manual (Becker et al. 2020). Analyses were performed on a continuous-flow QuAAtro39 AutoAnalyzer (Seal Analytical) with low detection limits (nitrate + nitrite [N+N]: 0.05 µM). Finally, primary productivity was estimated from uptake of 13C-bicarbonate in 24 h deck board incubation experiments following the methods of Hama et al. (1983) and Imai et al. (2002). Experiments were duplicated to compare production of <3 µm and >3 µm phytoplankton cells. For the <3 µm size fraction, samples were size-fractionated after incubation by filtering through 20 µm Nitex mesh followed by 3 µm pore-size polycarbonate membranes before collecting cells onto glass fiber filters (GFFs) (0.7 µm effective pore size). The other set of samples was collected directly onto GFFs. All GFFs were acid-fumed, dried, and analyzed for 13C/12C ratios and particulate organic carbon content at the UC Davis Stable Isotope Facility. C:chl conversion factors were determined with light microscope analysis and used to calculate phytoplankton carbon in µg C L^-1^ (Strom et al. 2016).

### Rhizaria sample analysis

Samples underwent a two-stage settling process in preparation for inverted light microscope analysis. Only one-fourth of the sample volume (100 mL) was analyzed. Each sample was settled for 48 h at 4°C in a 500 mL conical tube. 90 mL was removed with a peristaltic pump and the remaining 10 mL of settled material was carefully resuspended by mixing and transferred with a Pasteur pipette into a cylindrical chamber where it settled for 48 h at 4°C. To allow for nuclei visualization, 1 mL of 10 µg mL^-1^ DAPI stain was also added to the final sample volume during the second settling stage. Rhizarians were identified, counted, and measured with an eyepiece micrometer under light microscopy (Zeiss IM 35) at 200x and 320x.

Photographs were taken with a Google Pixel 6 camera (50 MP, f/1.9, 25 mm). Each individual was classified to at least phylum level (i.e. Foraminifera and Radiozoa also known as Radiolaria) or other major taxonomic group (i.e. Phaeodaria) based on the phylogeny of Cavalier-Smith (2018) and Nakamura et al. (2020) by referring to numerous visual guides (Nigrini and Moore, Jr. 1979, Kling and Boltovskoy 1999, Lee et al. 2000, Suzuki et al. 2009, Kimoto 2015, Nakamura and Suzuki 2015, Suzuki and Not 2015, Takagi et al. 2019, Munir et al. 2020, Mansour et al. 2021, Laget et al. 2023). Group-specific morphological and subcellular features were used for identification (Table 1). Subsequent lower taxonomic classifications were assigned where possible, including to species in two cases (*Sticholonche zanclea* and *Protocystis acornis*). Those individuals that could not be identified as belonging to a particular taxon but still displayed Rhizaria characteristics were classified as “Unknown Rhizaria”. Additionally, morphotypes were identified within each taxonomic group based on differences in appearance, morphology, and size of the endoplasm and spines. It is likely that some morphotypes represent different life stages or morphological variants of the same species.

**Table 1.**
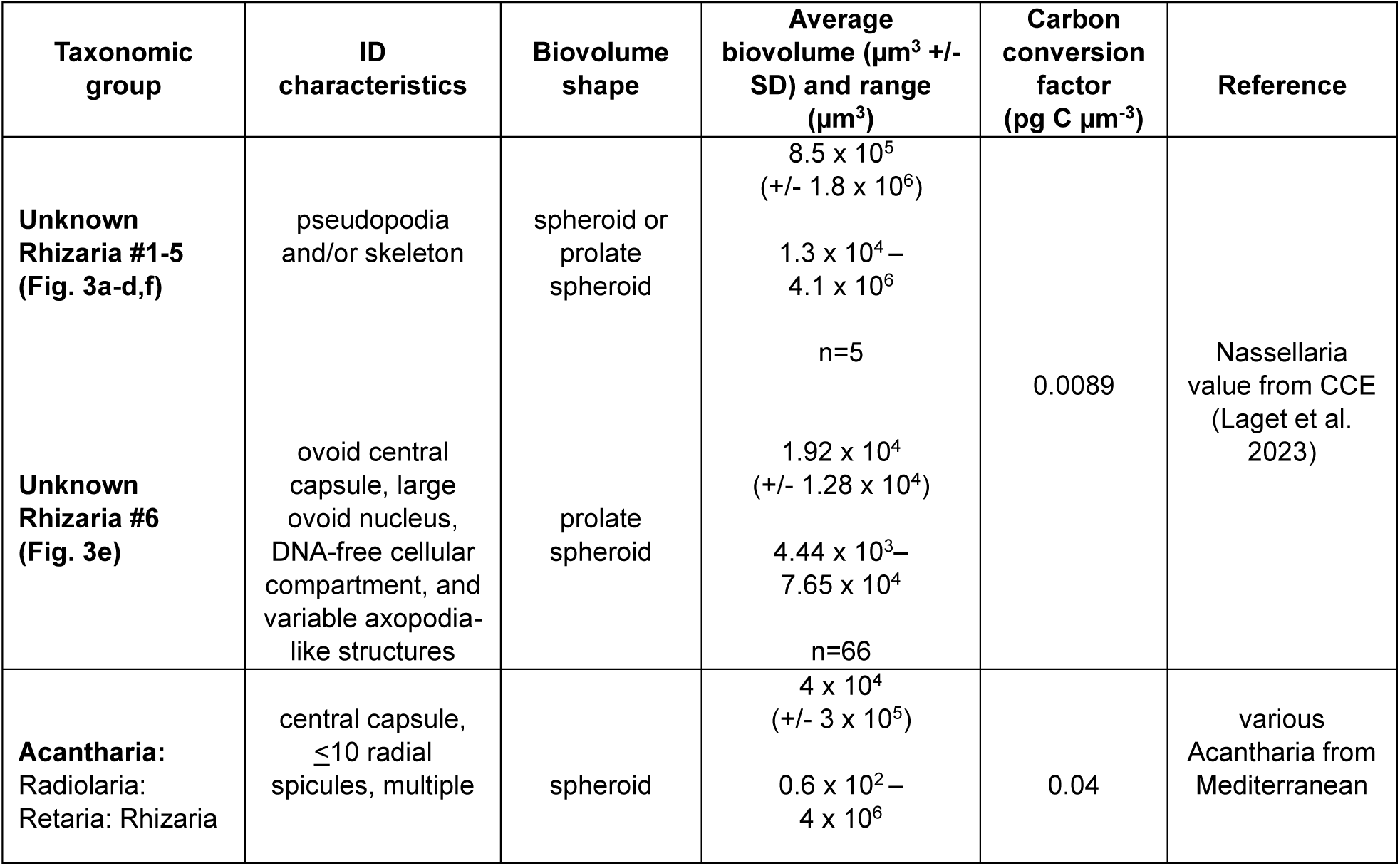
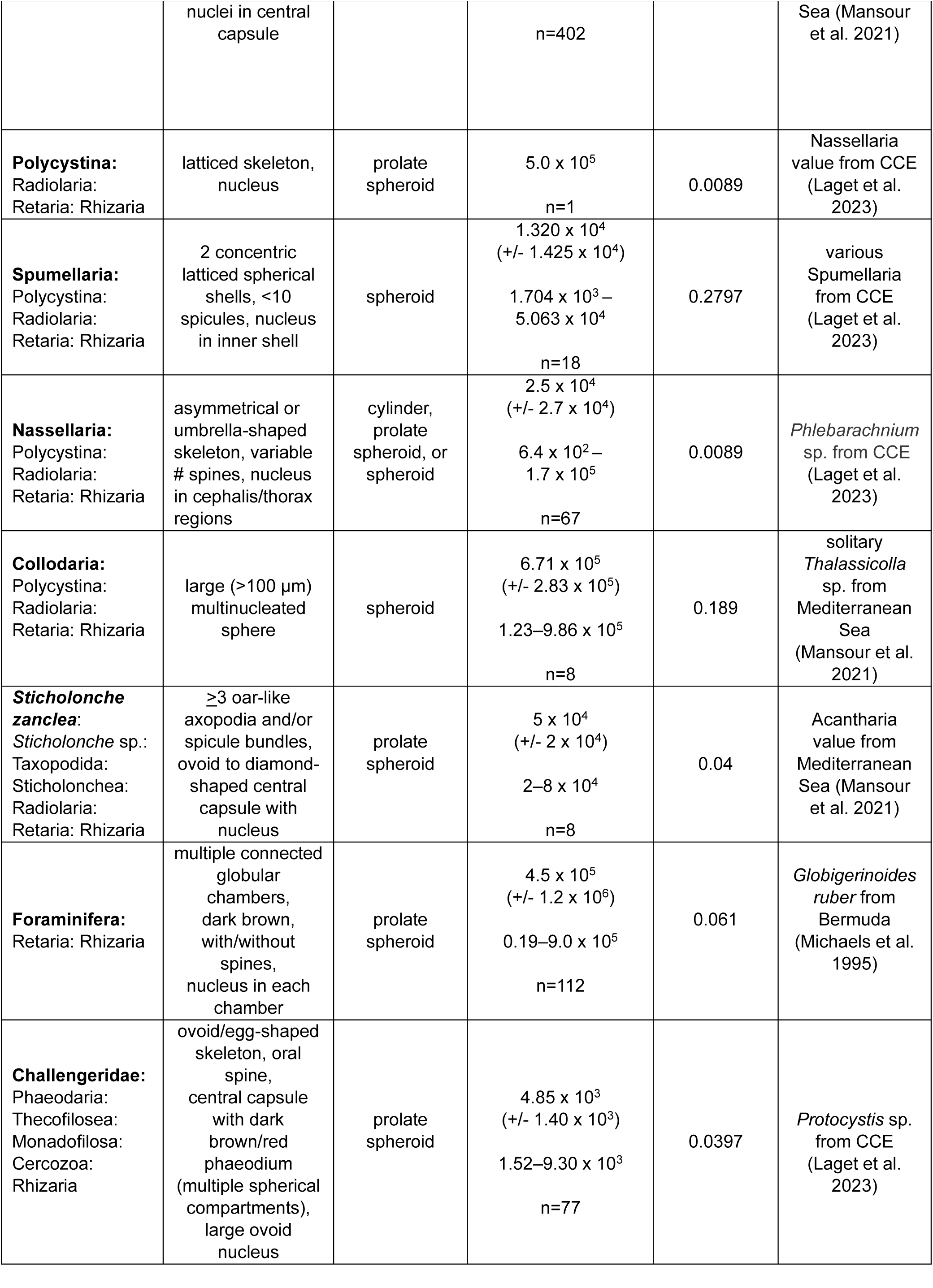
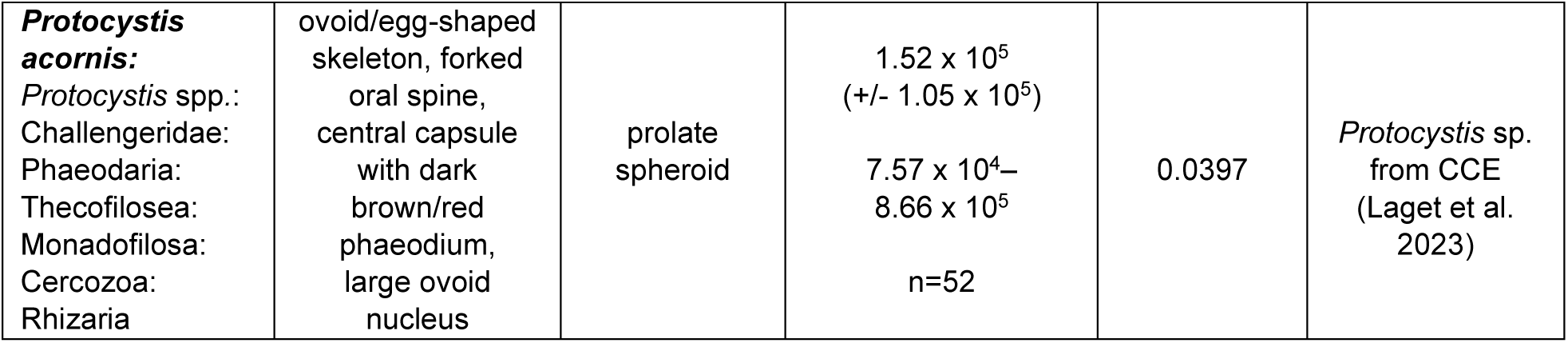
Criteria for Rhizaria identification and biomass estimation. *ID characteristics:* identifying characteristics used to classify taxa. *Biovolume shape:* 3D geometric shape assigned to each individual; corresponding volumetric equations are listed in Supplemental Table 2. *Average and range of biovolume:* listed for each taxonomic group; individuals lacking nuclei were not included in calculations. *Carbon conversion factor:* carbon density used for volume-to-biomass conversion for each taxonomic group. *Reference:* source of the carbon conversion factor; CCE=California Coastal Ecosystem.

Epifluorescence illumination utilizing UV and blue excitation wavelengths (to visualize blue-fluorescing DAPI-stained nuclei and yellow-to-red fluorescing photopigments, respectively) was used to confirm presence of protists alive at the time of fixation, including algal cells interacting with Rhizaria (Gowing 1989, Takahashi et al. 2003, Suzuki et al. 2009). Nuclei are located in the endoplasm of the central capsule of Radiolarians (Anderson 1983, Suzuki and Not 2015) and in the intra-shell cytoplasm of Foraminifera (Hemleben et al. 1989, Schiebel and Hemleben 2005); therefore, DAPI stain elucidated key aspects of cell structure and organization. Positive incidence of algal cell interaction was defined as widespread, yellow-to-red fluorescence that emanated from within the endoplasm of the central capsule for Radiolaria or chambers for Foraminifera. Algal cells interacting with Rhizarians also presented as small (< ∼5 µm) fluorescent puncta inside the central capsule or chambers. Hosts often exhibited both widespread fluorescence and more defined, bright puncta. In Acantharians, symbionts colonize either the extracapsulum (the cytoplasmic region outside the central capsule), or the intracapsulum (the endoplasm within the central capsule) and are contained in perialgal vacuoles (Anderson 1983, Suzuki and Not 2015, Decelle and Not 2015). Foraminifera also contain their symbionts in perialgal vacuoles, which are located in the cytoplasm-rhizopodial network and, through rhizopodial streaming by the host, can be moved inside the shell (Hemleben et al. 1989). In this study we could not determine whether the widespread autofluorescence or small puncta seen in Rhizarians represented truly symbiotic algal cells versus commensals or “hitchhikers”, so these incidences were defined as algal cell interactions.

Rhizarians with captured prey items were also identified. For Radiolaria and Foraminifera, positive incidence of prey capture was defined as an organism of an identifiable nature (e.g., diatom, tintinnid, etc.) with a nucleus and other intracellular material (i.e. not an empty diatom frustule) or, if unknown, with a nucleus and clear cell boundaries, that was visibly stuck through the Rhizarian’s spines/spicules (which are attached to the sticky, cytoplasmic pseudopodal network), and/or to the central capsule/shell. Additional evidence of prey capture was the presence of large (>5 µm) autofluorescent puncta or regions within the central capsule of Acantharians (only three cases) or within the phaeodium vacuoles of Challengeridae that indicated ingested algal prey. We recognize the presence of possible errors in prey capture assessment with our methods. At the least, we present quantitative and qualitative data to suggest the physical possibility of predator-prey interactions, even though predator-prey encounter rates may have been enhanced by our multi-step concentration and settlement sample preparation process.

### Biovolume measurements and biomass calculations

Geometric equations from Hillebrand et al. (1999) were used to calculate biovolumes of Rhizarians that had nuclei, based on an assigned shape (Table 1, with further details in Supplemental Table 2). Nuclei were observed in most individuals within each taxa. Individuals without DAPI-staining nuclei were omitted from biovolume and biomass analyses because these were assumed to be empty skeletons or dead cells. Where applicable, ectoplasm (usually deteriorated or not visible), pseudopodia/axopodia extensions, and spicules/spines were not included in biovolume measurements; this is now common practice (Stukel et al. 2018, Ikenoue et al. 2019, Mansour et al. 2021). We know of only three reports of Rhizaria cell carbon density: Michaels et al. (1995), Mansour et al. (2021), and Laget et al. (2023). These studies collected Rhizarian specimens of varying taxonomic groups from different marine systems and include at least one case of small sample size. Nonetheless, carbon density values determined by these researchers were used to calculate the biomass of each individual in this study (Table 1). The most relevant conversion factors were chosen for each group, based either on sample collection location or morphological/taxonomic similarity. Mansour et al. (2021) compiled various Mediterranean Sea and Southern Ocean Nassellarians and obtained a carbon density value ∼150x greater than that of Laget et al. (2023) (1.472 vs. 0.0089 pg C µm^-3^, respectively). Laget et al. (2023) collected “large” *Phlebarachnium* sp. Nassellarians. Even though we did not identify any Nassellarians that resembled this genus’ morphology, 0.0089 pg C µm^-3^ was used because not only is it a small value that resulted in conservative biomass estimates for this taxon, but the samples used in that analysis were from coastal waters in the California Current Ecosystem (CCE), a system co-located with the NGA in the eastern Pacific Ocean. This Nassellaria conversion factor was also used in biomass calculations for Unknown Rhizaria because it was the lowest carbon density out of the suite used in this study, yielding conservative biomass estimates for these individuals. The carbon density of *Protocystis* sp. collected from the CCE was used to calculate biomass of Challengeridae and *Protocystis acornis.* Finally, the Acantharia carbon density value was used to calculate biomass of *Sticholonche zanclea* (Taxopodida) cells because the groups share similar morphologies.

### Community analysis

The relationship between Rhizaria communities at different stations and depths was assessed using the vegan community ecology package (v2.6.6.1; Oksanen et al. 2024) performed in R Statistical Software (v4.4.1; R Core Team 2024). Communities were compared using abundance data in a two-dimension non-metric multidimensional scaling ordination (“metaMDS” function) with the Bray-Curtis dissimilarity index (stress=0.112, non-metric fit R^2^=0.987).

## RESULTS

### Oceanographic setting

During Summer 2023 in the NGA, a pronounced density gradient was evident nearshore in the upper 30 m (GAK1, Figure 2a). The pycnocline was deeper farther offshore at GAK9 and GAK15 (∼18-27 m, Figure 2a) compared to ∼5 m on the inner to mid shelf (GAK1 and GAK5, Figure 2a). Salinity ranged from ∼32.0 to 32.3 at the surface at GAK5, GAK9, and GAK15, with a prominent low-salinity, freshwater signature at GAK1 (∼27.7, Figure 2b). Temperature at the surface was lowest at GAK1 (9.5°C, Figure 2c) and increased with distance offshore. The cooler inshore temperatures and low salinities were likely influenced by coastal freshwater inputs transported westward by the Alaska Coastal Current (Figure 1). Nitrate increased with distance offshore on the Seward line and was greatest at GAK15 (Figure 2d), where dissolved Fe (dFe) was lowest (<0.2 nM, Figure 2e). Biological indicators were also examined, including size-fractionated chlorophyll-a (large cells >20 µm and small cells <20 µm that represent potential Rhizaria algal prey and symbionts/commensals, respectively) and primary productivity. Large cell chlorophyll-a maxima were located between 10 and 40 m; the highest was at GAK1 (10 m, 0.7 mg L^-1^, Figure 2f). Small cell chlorophyll-a maxima were at 10 m and similar in magnitude (∼0.9 mg L^-1^) at on-shelf stations, whereas offshore concentrations were highest at 30-40 m (e.g. 0.3 mg L^-1^ at GAK15; Figure 2g). Total chlorophyll-a and dFe were lowest at GAK15; coupled with elevated nitrate, these properties indicate that high-nitrate low-chlorophyll (HNLC) conditions indicative of Fe limitation were present in offshore NGA waters during summer 2023. Integrated primary production across Seward line stations GAK1, GAK5, GAK9, and GAK15 averaged 935 mg C m^-2^ d ^-1^ and was greatest inshore (Figure 2h). Picophytoplankton (<3 µm) contributed over 50% of primary production at GAK5 and GAK9 (Figure 2h).

**Figure 2.**
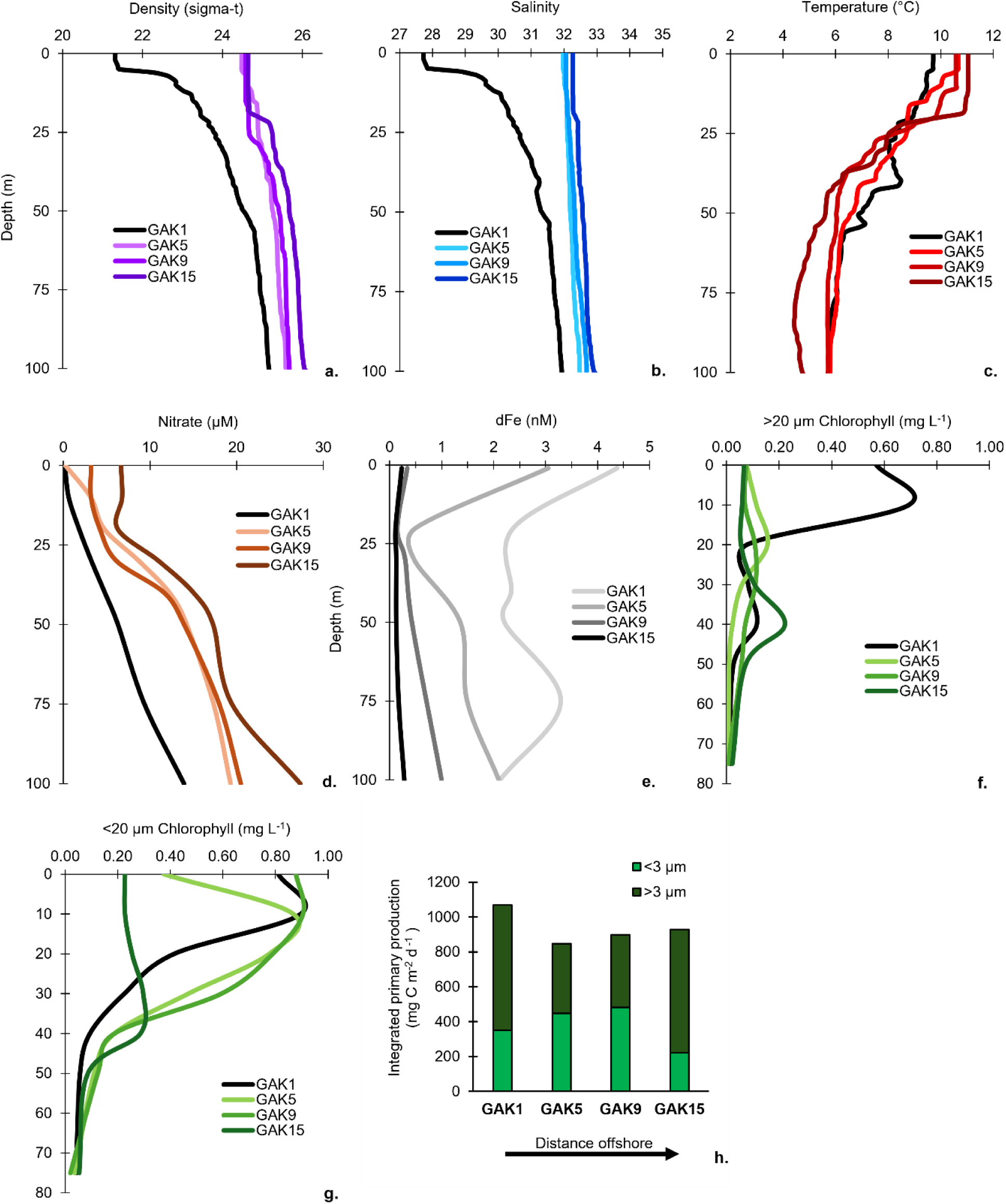
Vertical profiles of oceanographic parameters and cross-shelf primary productivity at GAK1, GAK5, GAK9, and GAK15 in summer 2023. a) density (sigma-theta) at 0-100 m, b) salinity at 0-100 m, c) temperature (°C) at 0-100 m, d) nitrate (µM) at 0-100 m, e) dissolved Fe, f) >20 µm chlorophyll-a (µg L^-1^) at 0-75 m, g) <20 µm chlorophyll-a (µg L^-1^) at 0-75 m, and h) integrated primary productivity (mg C m^2^ d^-1^) from cells separated into <3 µm and >3 µm size-fractions.

### Rhizaria diversity

Rhizaria were classified into ten taxonomic groups in this study (Table 1). As described in Methods, Rhizarians were assigned to the lowest taxonomic classification possible based solely on morphology as visualized through an inverted epi-fluorescence microscope. Some individuals exhibited Rhizaria characteristics but were unidentifiable and therefore could only be classified as “Unknown Rhizaria”, of which there were six species/morphotypes (Figure 3). One Unknown Rhizaria morphotype did not closely resemble any organism in the literature but was abundant in the NGA (Unknown Rhizaria #6, Figure 3e; max abundance 6.4 cells L^-1^, Table 2). Rhizaria cells ranged in volume from 0.6 x 10^2^ µm^3^ (Acantharia) to 9.0 x 10^5^ µm^3^ (Foraminifera) (Table 2). Acantharia were variable in size (0.6 x 10^2^ to 4 x 10^6^ µm^3^; Table 1) and occurred in 20 different morphotypes (Figure 4a-t). These Radiolarians were multinucleated as seen previously by Suzuki et al. (2009). Individual *Sticholonche zanclea* from the single-species order Taxopodida (Figure 5a) varied between 2 and 8 x 10^4^ µm^3^ (Table 1). Nassellaria exhibited the greatest morphological diversity of the Polycystines, with 24 morphotypes (Figure 6e-z, iii-iv) that ranged in size from 6.4 x 10^2^ to 1.7 x 10^5^ µm^3^ (Table 1). Four Spumellaria (Figure 6a-d), one Collodaria (Figure 6ii), and two Unknown Polycystine (Figure 6v-vi) morphotypes were also identified. Five morphotypes of Foraminiferans were categorized based on general size (small, large, very large) and presence/absence of spines (Figure 7a-e). These multichambered, globular protists were by far the largest Rhizarians observed, varying in size from 0.19 to 9.0 x 10^5^ µm^3^ (Table 2). Four morphotypes of Challengeridae were found (Figure 8a-d), including individuals that displayed *Protocystis acornis* characteristics, mainly a forked oral spine (Figure 8d). Challengeridae (including *P. acornis*) volumes varied between 1.52 x 10^3^ and 8.66 x 10^5^ µm^3^ (Table 1).

**Figure 3.**
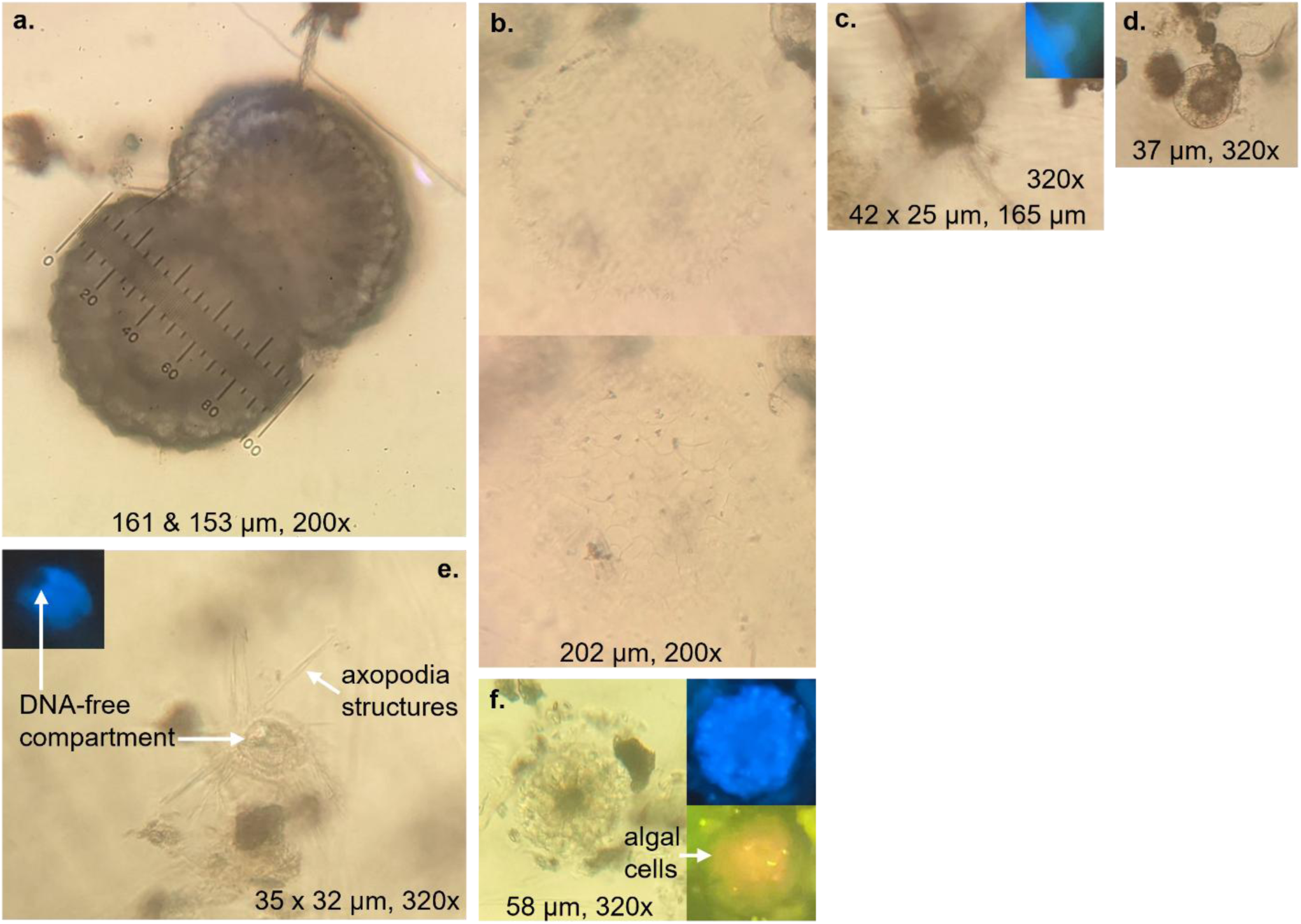
Unknown Rhizaria diversity. Dimensions of the central capsule(s) used in biovolume calculations, spicule extent length (**d** and **f**), and magnification are noted. Inset images using UV and blue light illumination show DAPI-stained nuclei and fluorescent photopigments, respectively. **a)** two large cells with scalloped outer boundaries. **b)** two views of a large, empty, latticed sphere with short spines. **c)** irregular shaped cell with spherical endoplasm and at least 8 spines. **d)** small sphere with large, dense endoplasm. **e) Unknown Rhizaria #6:** cell with an ovoid central capsule, a large ovoid nucleus, a DNA-free compartment, and variable axopodia-like structures. **f)** small sphere with small dense endoplasm and at least 4 spines.

**Figure 4.**
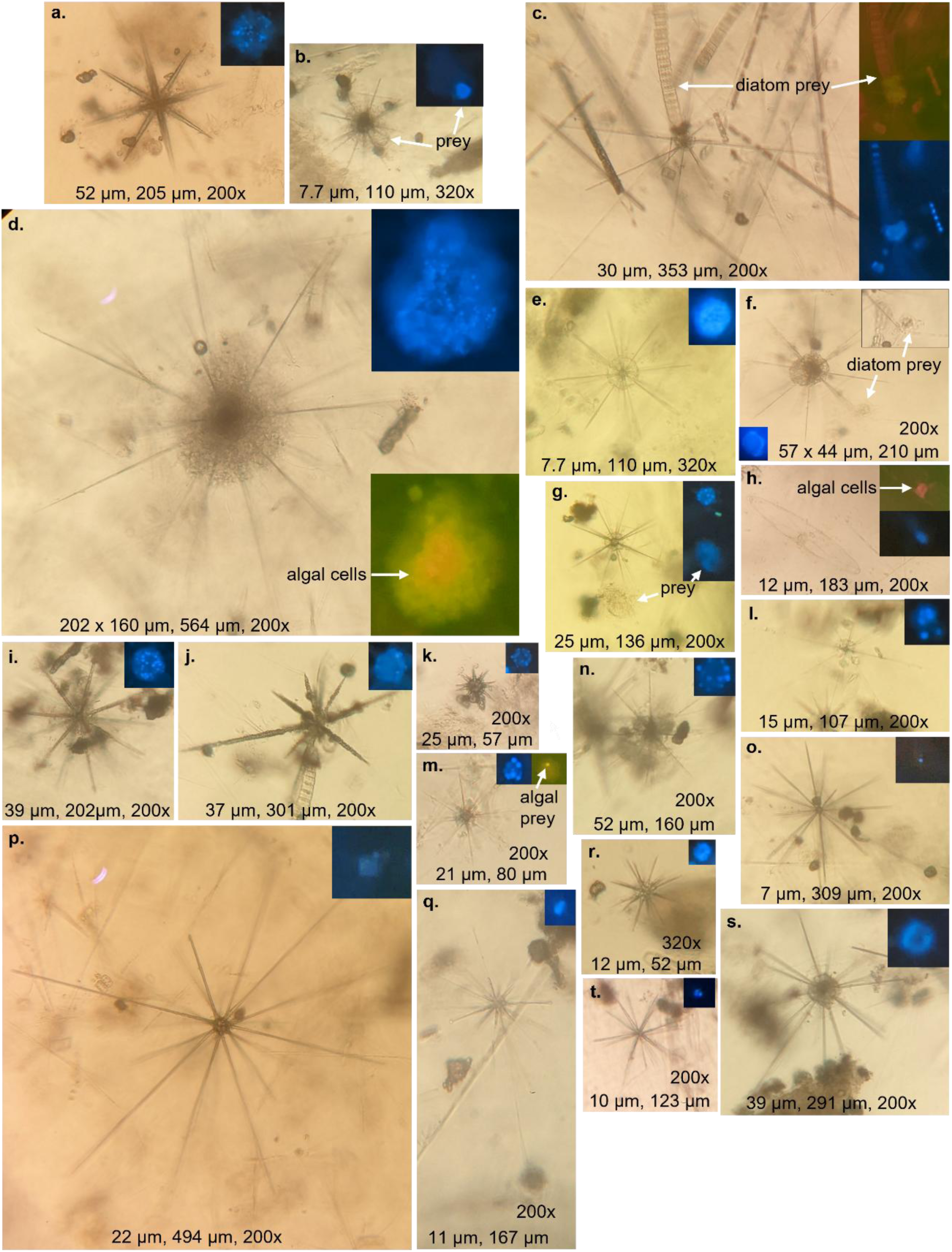
Acantharia diversity. Dimensions of the central capsule used in biovolume calculations, spicule extent length, and magnification are noted. Inset images using UV and blue light illumination show DAPI-stained nuclei and fluorescent photopigments, respectively. **a-t)** Acantharia morphotypes: **b)** and **g)** with captured unknown prey containing nuclei, **c)** and **f)** with captured diatom prey, **d)** and **h)** with intracapsulum-associated red-orange and red algal cells, and **m)** with algal prey in endoplasm.

**Figure 5.**
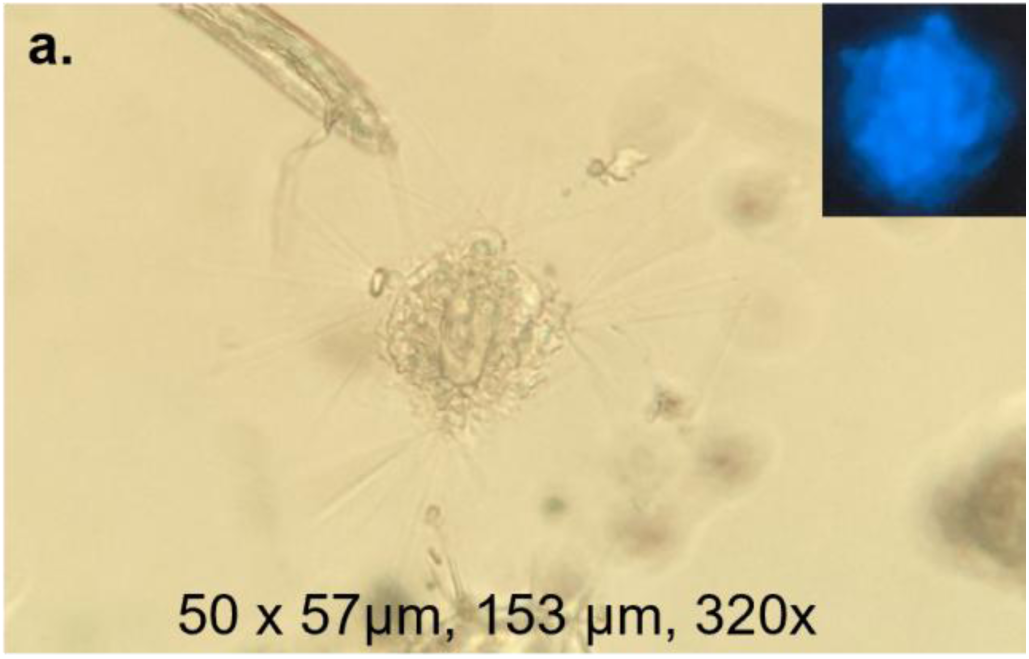
Taxopodida diversity. Dimensions of the central capsule used in biovolume calculations, spicule extent length, and magnification are noted. Inset images using UV and blue light illumination show DAPI-stained nuclei and fluorescent photopigments, respectively. a) *Sticholonche zanclea*.

**Figure 6.**
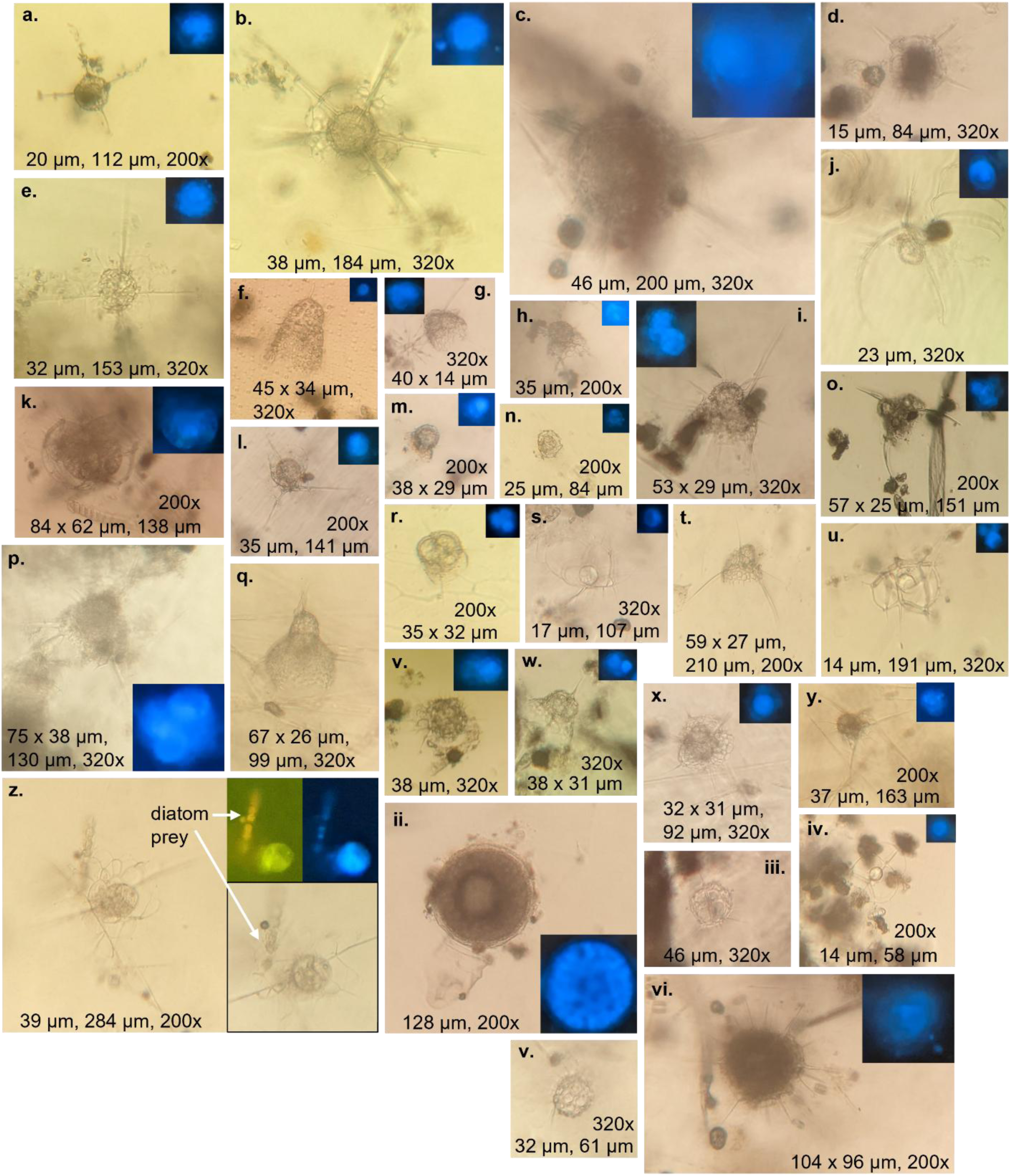
Polycystina diversity. Dimensions used in biovolume calculations, spine extent length, and magnification are noted. Inset images using UV and blue light illumination show DAPI-stained nuclei and fluorescent photopigments, respectively. a-d) Spumellaria morphotypes. e-z, iii-iv) Nassellaria morphotypes: z) with diatom prey stuck to skeleton. ii) Collodaria morphotype. v-vi) Unknown Polycystina morphotypes.

**Figure 7.**
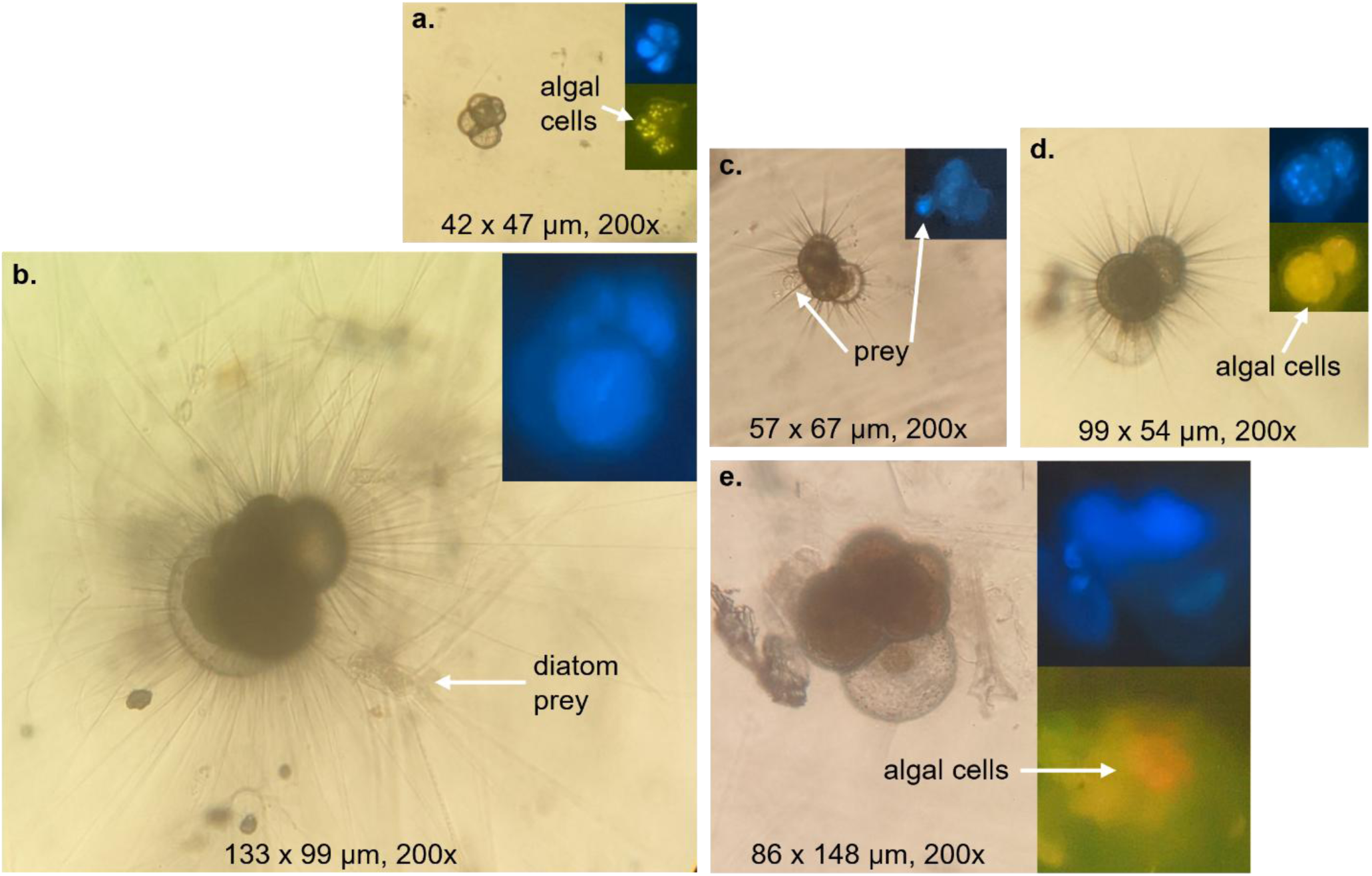
Foraminifera diversity. Dimensions used in biovolume calculations and magnification are noted. Inset images using UV and blue light illumination show DAPI-stained nuclei and fluorescent photopigments, respectively. a-e) Foraminifera morphotypes: a) small (<90 µm), non-spinose (note intrashell-associated yellow algal cells in this individual), b) very large (>130 µm), spinose; *Chaetoceros* spp. prey captured in spines, c) small (<90 µm), spinose; unknown prey captured in spines and to chamber, d) large (>90 µm), spinose (note intrashell-associated golden-yellow algal cells in this individual), and e) large (>90 µm), non-spinose (note intrashell-associated red-orange algal cells in this individual).

**Figure 8.**
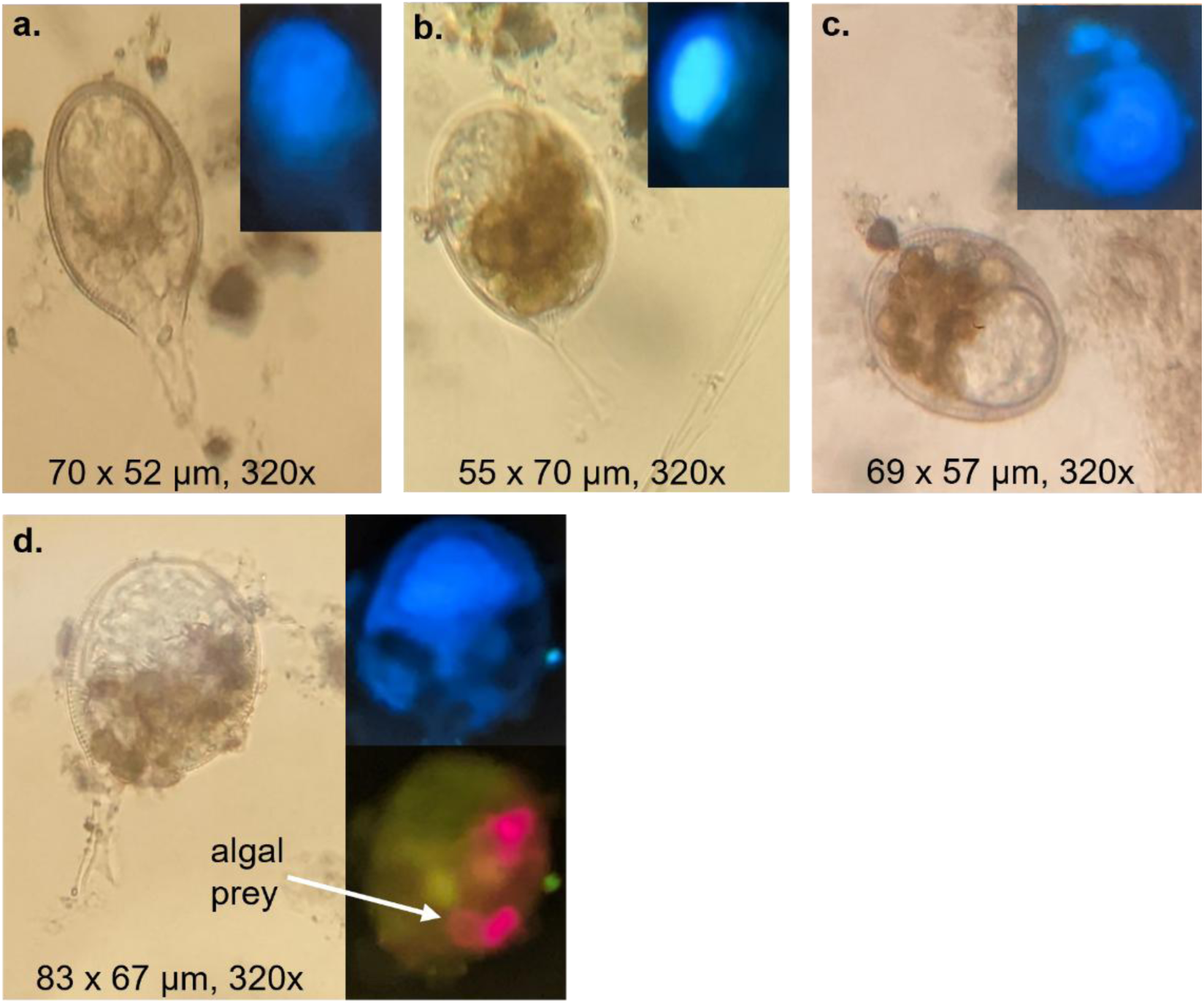
Phaeodaria diversity. Dimensions used in biovolume calculations and magnification are noted. Inset images using UV and blue light illumination show DAPI-stained nuclei and fluorescent photopigments, respectively. **a-c)** Challengeridae morphotypes: **a)** with a pointed oral spine, **b)** with an oral spine of uncertain shape, and **c)** no oral spine seen. **d)** *Protocystis acornis* with digested algal prey in phaeodium.

**Table 2.** Taxon-specific abundances (cells L ) at each station and depth interval, followed by (# observed).

|  | GAK1<br>0-20 m | GAK1<br>30-50 m | GAK1<br>60-80 m | GAK1<br>100-250 m | GAK5<br>0-20 m | GAK5<br>30-50 m | GAK5<br>60-80 m | GAK5<br>90-150 m | GAK9<br>0-20 m | GAK9<br>30-50 m | GAK9<br>60-80 m | GAK9<br>100-250 m | GAK15<br>0-20 m | GAK15<br>30-50 m | GAK15<br>60-100 m | GAK15<br>500-1000 m | Total |
| --- | --- | --- | --- | --- | --- | --- | --- | --- | --- | --- | --- | --- | --- | --- | --- | --- | --- |
| Unknown<br>Rhizaria #1-5 | 0 (0) | 0 (0) | 0 (0) | 0 (0) | 0 (0) | 0.45 (4) | 0 (0) | 0 (0) | 0 (0) | 0 (0) | 0.11 (1) | 0.11 (1) | 0 (0) | 0 (0) | 0.11 (1) | 0 (0) | 0.78 (7) |
| Unknown<br>Rhizaria #6 | 0 (0) | 0 (0) | 0 (0) | 0 (0) | 0 (0) | 0 (0) | 0 (0) | 0 (0) | 0 (0) | 0.23 (2) | 6.4 (56) | 0 (0) | 0 (0) | 0 (0) | 0.68 (6) | 0.23 (2) | 7.5 (66) |
| Acantharia | 0 (0) | 0 (0) | 0 (0) | 0.68 (6) | 0.57 (5) | 0 (0) | 3.4 (30) | 5.3 (47) | 0.11 (1) | 5.8 (51) | 11 (100) | 5.2 (46) | 0.34 (3) | 1.4 (12) | 12 (109) | 2.4 (21) | 48 (431) |
| Polycystina | 0 (0) | 0 (0) | 0 (0) | 0 (0) | 0 (0) | 0 (0) | 0 (0) | 0 (0) | 0 (0) | 0 (0) | 0 (0) | 0 (0) | 0 (0) | 0.11 (1) | 0 (0) | 0.11 (1) | 0.22 (2) |
| Spumellaria | 0 (0) | 0 (0) | 0 (0) | 0 (0) | 0 (0) | 0.91 (8) | 0.45 (4) | 0.11 (1) | 0 (0) | 0.45 (4) | 0.23 (2) | 0 (0) | 0 (0) | 0 (0) | 0 (0) | 0 (0) | 2.2 (19) |
| Nassellaria | 0 (0) | 0 (0) | 0 (0) | 0 (0) | 0 (0) | 0.57 (5) | 0.57 (5) | 0.68 (6) | 0 (0) | 0.23 (2) | 1.5 (13) | 1.4 (12) | 0 (0) | 0.68 (6) | 2.5 (22) | 1.0 (9) | 9.1 (80) |
| Collodaria | 0 (0) | 0 (0) | 0 (0) | 0 (0) | 0 (0) | 0 (0) | 0 (0) | 0 (0) | 0 (0) | 0 (0) | 0 (0) | 0.91 (8) | 0 (0) | 0 (0) | 0 (0) | 0 (0) | 0.91 (8) |
| Sticholonche<br>zanclea | 0 (0) | 0 (0) | 0 (0) | 0 (0) | 0 (0) | 0.11 (1) | 0 (0) | 0.11 (1) | 0 (0) | 0.23 (2) | 0 (0) | 0.23 (2) | 0 (0) | 0 (0) | 0.34 (3) | 0 (0) | 1.0 (9) |
| Foraminifera | 0 (0) | 0 (0) | 0.23 (2) | 0 (0) | 2.6 (23) | 0.79 (7) | 0.34 (3) | 0.23 (2) | 1.8 (16) | 2.5 (22) | 0.57 (5) | 0.45 (4) | 4.1 (36) | 1.9 (17) | 0.34 (3) | 0 (0) | 16 (140) |
| Challengeridae | 0 (0) | 0 (0) | 0 (0) | 0 (0) | 0 (0) | 0 (0) | 0.23 (2) | 2.5 (22) | 0.11 (1) | 0.23 (2) | 3.1 (27) | 1.6 (14) | 0 (0) | 0 (0) | 1.7 (15) | 0 (0) | 9.5 (83) |
| Protocystis<br>acornis | 0 (0) | 0 (0) | 0 (0) | 0 (0) | 0 (0) | 0 (0) | 0 (0) | 1.2 (11) | 0 (0) | 0 (0) | 1.9 (17) | 2.0 (18) | 0 (0) | 0 (0) | 0.68 (6) | 0 (0) | 5.8 (52) |

### Rhizaria ecology

#### Distributions and community composition

In Summer 2023, Rhizaria abundances generally increased with depth and with distance offshore. On the inner shelf at GAK1 and GAK5, abundances were greatest in the deep water (<1.0 and 10 cells L^-1^, respectively, Figure 9a). Abundances reached their depth maxima in intermediate waters both on the outer shelf and in the open ocean (GAK9 and GAK15, 25 and 19 cells L^-1^, respectively, Figure 9a). Across the NGA, Rhizaria abundances were generally lowest at the surface, ranging from 0 to 4 cells L^-1^ (Figure 9a). In fact, no Rhizarians were detected at the nearshore GAK1 surface and mid depths. At GAK9 and GAK15, deep-water abundances diminished at least twofold relative to the intermediate water quantities. In contrast to abundance, Rhizaria biomass did not show distinct cross-shelf or vertical patterns, since each station had a different depth maximum (Figure 9b). At GAK5, biomass peaked at the surface (105 ng C L^-1^, Figure 9b) where it comprised mainly large Foraminiferans (100 ng C L^-1^, Table 3). At GAK9, biomass was highest in the deep water (143 ng C L^-1^, Figure 9b) with a large proportion of Collodarians (115 ng C L^-1^, Table 3). Off the shelf at GAK15, biomass was greatest at mid depths (137 ng C L^-1^, Figure 9b).

**Figure 9.**
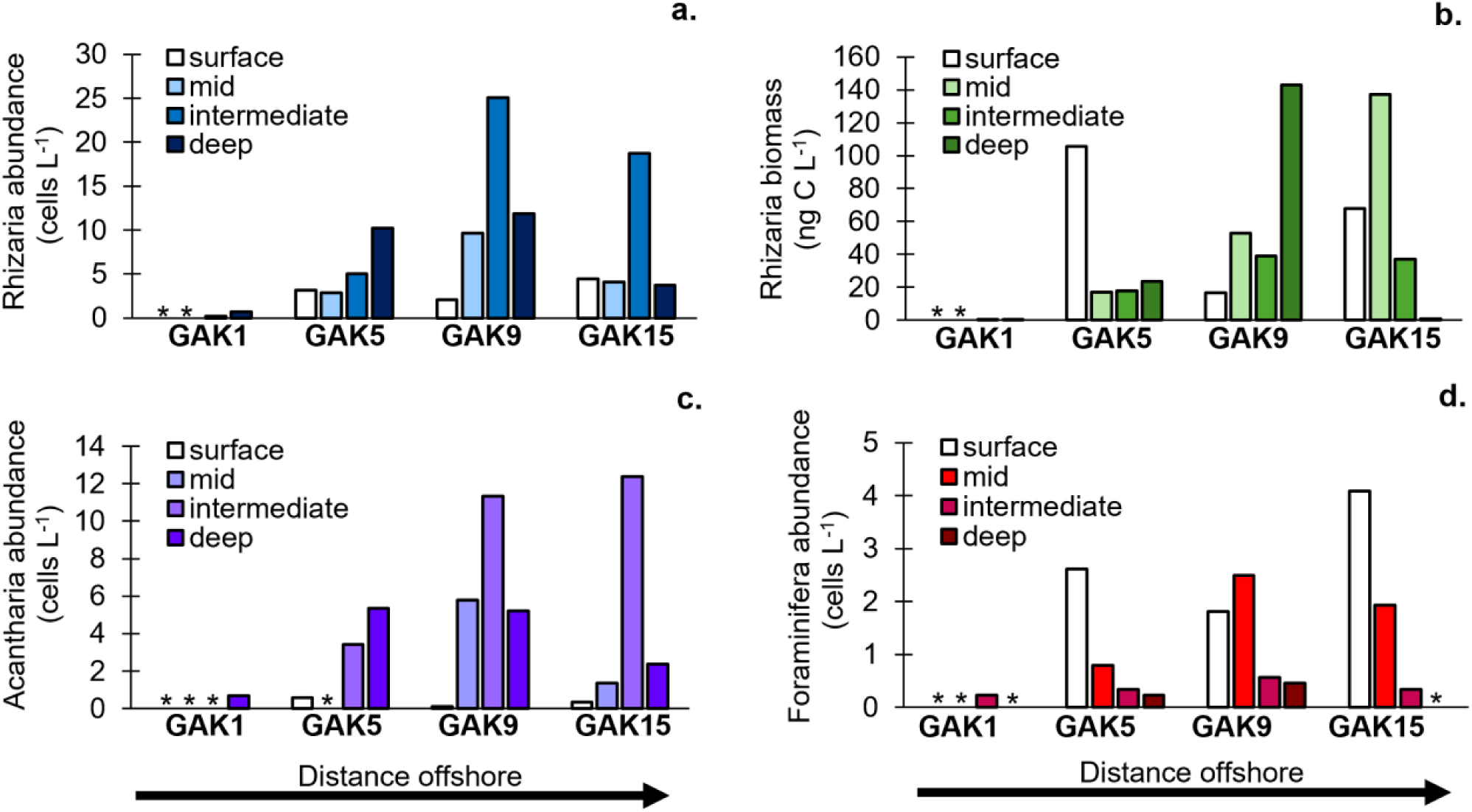
Distributions of Rhizaria. a) abundance (cells L^-1^) (n=897), b) biomass (ng C L^-1^) (n=819), c) Acantharia abundance (cells L^-1^) (n=431) and d) Foraminifera abundance (cells L^-1^) (n=140) at different depth intervals (see Supp. Table 1) across the Seward line. Depth intervals are shown in Table 1. * = none detected.

**Table 3.**
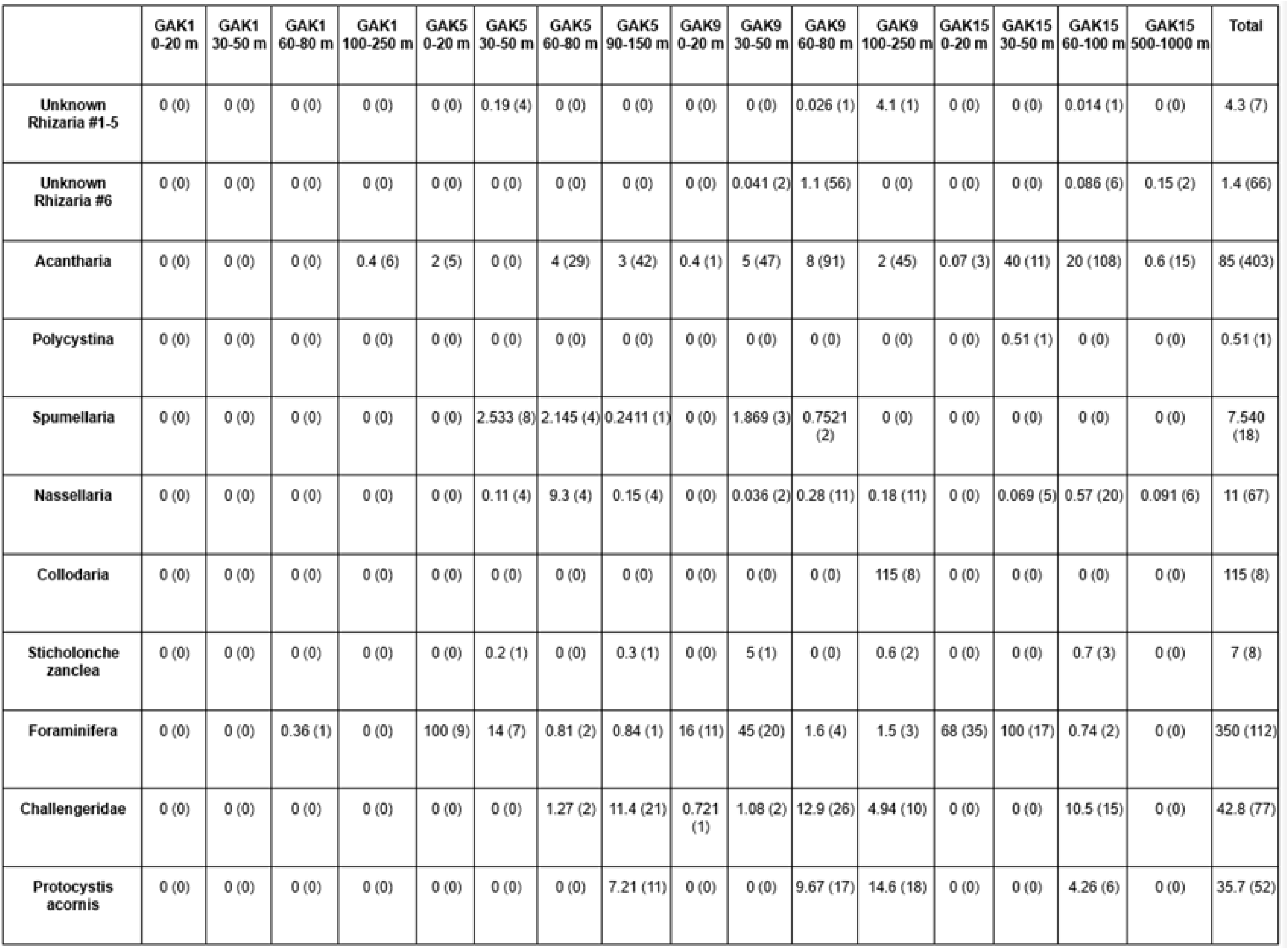
Taxon-specific biomass for each station and depth interval (ng C L-1) followed by (sample size).

|  | GAK1<br>0-20 m | GAK1<br>30-50 m | GAK1<br>60-80 m | GAK1<br>100-250 m | GAK5<br>0-20 m | GAK5<br>30-50 m | GAK5<br>60-80 m | GAK5<br>90-150 m | GAK9<br>0-20 m | GAK9<br>30-50 m | GAK9<br>60-80 m | GAK9<br>100-250 m | GAK15<br>0-20 m | GAK15<br>30-50 m | GAK15<br>60-100 m | GAK15<br>500-1000 m | Total |
| --- | --- | --- | --- | --- | --- | --- | --- | --- | --- | --- | --- | --- | --- | --- | --- | --- | --- |
| Unknown<br>Rhizaria #1-5 | 0 (0) | 0 (0) | 0 (0) | 0 (0) | 0 (0) | 0.19 (4) | 0 (0) | 0 (0) | 0 (0) | 0 (0) | 0.026 (1) | 4.1 (1) | 0 (0) | 0 (0) | 0.014 (1) | 0 (0) | 4.3 (7) |
| Unknown<br>Rhizaria #6 | 0 (0) | 0 (0) | 0 (0) | 0 (0) | 0 (0) | 0 (0) | 0 (0) | 0 (0) | 0 (0) | 0.041 (2) | 1.1 (56) | 0 (0) | 0 (0) | 0 (0) | 0.086 (6) | 0.15 (2) | 1.4 (66) |
| Acantharia | 0 (0) | 0 (0) | 0 (0) | 0.4 (6) | 2 (5) | 0 (0) | 4 (29) | 3 (42) | 0.4 (1) | 5 (47) | 8 (91) | 2 (45) | 0.07 (3) | 40 (11) | 20 (108) | 0.6 (15) | 85 (403) |
| Polycystina | 0 (0) | 0 (0) | 0 (0) | 0 (0) | 0 (0) | 0 (0) | 0 (0) | 0 (0) | 0 (0) | 0 (0) | 0 (0) | 0 (0) | 0 (0) | 0.51 (1) | 0 (0) | 0 (0) | 0.51 (1) |
| Spumellaria | 0 (0) | 0 (0) | 0 (0) | 0 (0) | 0 (0) | 2.533 (8) | 2.145 (4) | 0.2411 (1) | 0 (0) | 1.869 (3) | 0.7521 (2) | 0 (0) | 0 (0) | 0 (0) | 0 (0) | 0 (0) | 7.540 (18) |
| Nassellaria | 0 (0) | 0 (0) | 0 (0) | 0 (0) | 0 (0) | 0.11 (4) | 9.3 (4) | 0.15 (4) | 0 (0) | 0.036 (2) | 0.28 (11) | 0.18 (11) | 0 (0) | 0.069 (5) | 0.57 (20) | 0.091 (6) | 11 (67) |
| Collodaria | 0 (0) | 0 (0) | 0 (0) | 0 (0) | 0 (0) | 0 (0) | 0 (0) | 0 (0) | 0 (0) | 0 (0) | 0 (0) | 115 (8) | 0 (0) | 0 (0) | 0 (0) | 0 (0) | 115 (8) |
| Sticholonche<br>zanclea | 0 (0) | 0 (0) | 0 (0) | 0 (0) | 0 (0) | 0.2 (1) | 0 (0) | 0.3 (1) | 0 (0) | 5 (1) | 0 (0) | 0.6 (2) | 0 (0) | 0 (0) | 0.7 (3) | 0 (0) | 7 (8) |
| Foraminifera | 0 (0) | 0 (0) | 0.36 (1) | 0 (0) | 100 (9) | 14 (7) | 0.81 (2) | 0.84 (1) | 16 (11) | 45 (20) | 1.6 (4) | 1.5 (3) | 68 (35) | 100 (17) | 0.74 (2) | 0 (0) | 350 (112) |
| Challengeridae | 0 (0) | 0 (0) | 0 (0) | 0 (0) | 0 (0) | 0 (0) | 1.27 (2) | 11.4 (21) | 0.721 (1) | 1.08 (2) | 12.9 (26) | 4.94 (10) | 0 (0) | 0 (0) | 10.5 (15) | 0 (0) | 42.8 (77) |
| Protocystis<br>acornis | 0 (0) | 0 (0) | 0 (0) | 0 (0) | 0 (0) | 0 (0) | 0 (0) | 7.21 (11) | 0 (0) | 0 (0) | 9.67 (17) | 14.6 (18) | 0 (0) | 0 (0) | 4.26 (6) | 0 (0) | 35.7 (52) |

Acantharia was the most abundant taxon in the NGA followed by Foraminifera, which were about threefold less abundant (Figure 9c,d). Similar to the Rhizaria cross-shelf gradient, Acantharia and Foraminifera generally increased with distance offshore, while the vertical distributions exhibited opposite trends. Acantharia were more likely to be found below the surface, peaking at intermediate or deep depths depending on the location (maximum abundance 12 cells L^-1^ in GAK15 intermediate waters, Figure 9c). In contrast, most Foraminifera resided at or just below the surface, reaching 4.1 cells L^-1^ at GAK15 (Figure 9d).

Rhizaria community composition was evaluated in terms of three major taxonomic groups (Foraminifera, Radiolaria, and Phaeodaria) and by Radiolaria subgroup since it was the most diverse phylum. Foraminifera was consistently the dominant phylum in surface waters extending from the mid-shelf (GAK5) to the open ocean (GAK15) (82 to 92% relative abundance, Figure 10a). The exception was GAK1, which did not have any Rhizarians at the surface and mid depths. Foraminifera was the only phylum present at GAK1 intermediate depths (100% relative abundance, Figure 10a). Across all stations, Foraminifera influenced Rhizaria biomass to the greatest extent at 0-20 m and 30-50 m depths (73 to 100% relative biomass, Figure 10b) due to their large sizes. Due to the high Acantharia abundances, Radiolaria was the most abundant phylum across the NGA with >50% relative abundance at all station depths below the surface (excluding GAK1 60-80 m; Figure 10a). Acantharia comprised >60% of the Radiolarian community at all station depths except GAK5 30-50 m where they were absent (Figure 10c). Nassellaria was the second most abundant Radiolarian (range 3.4 to 36% relative abundance, Figure 10c) followed by Spumellaria (range 2.0 to 57% relative abundance, Figure 10c). Acantharia also dominated the Radiolarian biomass at most depths and locations (>70% relative biomass, Figure 10d) except for discrete samples (i.e. GAK5 30-50 m, 60-80 m and GAK15 100-250 m) where Spumellaria, Nassellaria, and Collodaria contributed higher biomass, respectively. Phaeodaria was the least abundant major clade overall; this group contributed anywhere from 2.4 to 37% relative abundance (Figure 10a) and was more likely to reside in intermediate and deep waters. This group comprised the majority of Rhizarian biomass at GAK5 90-150 m and GAK9 60-80 m (79 and 65% relative biomass, respectively, Figure 10b) where Radiolaria dominated in terms of abundance. This disproportionate contribution to biomass is because *P. acornis* individuals were on average one order of magnitude larger than Acantharia (1.52 x 10^5^ and 4 x 10^4^ µm^3^, respectively, Table 1).

**Figure 10.**
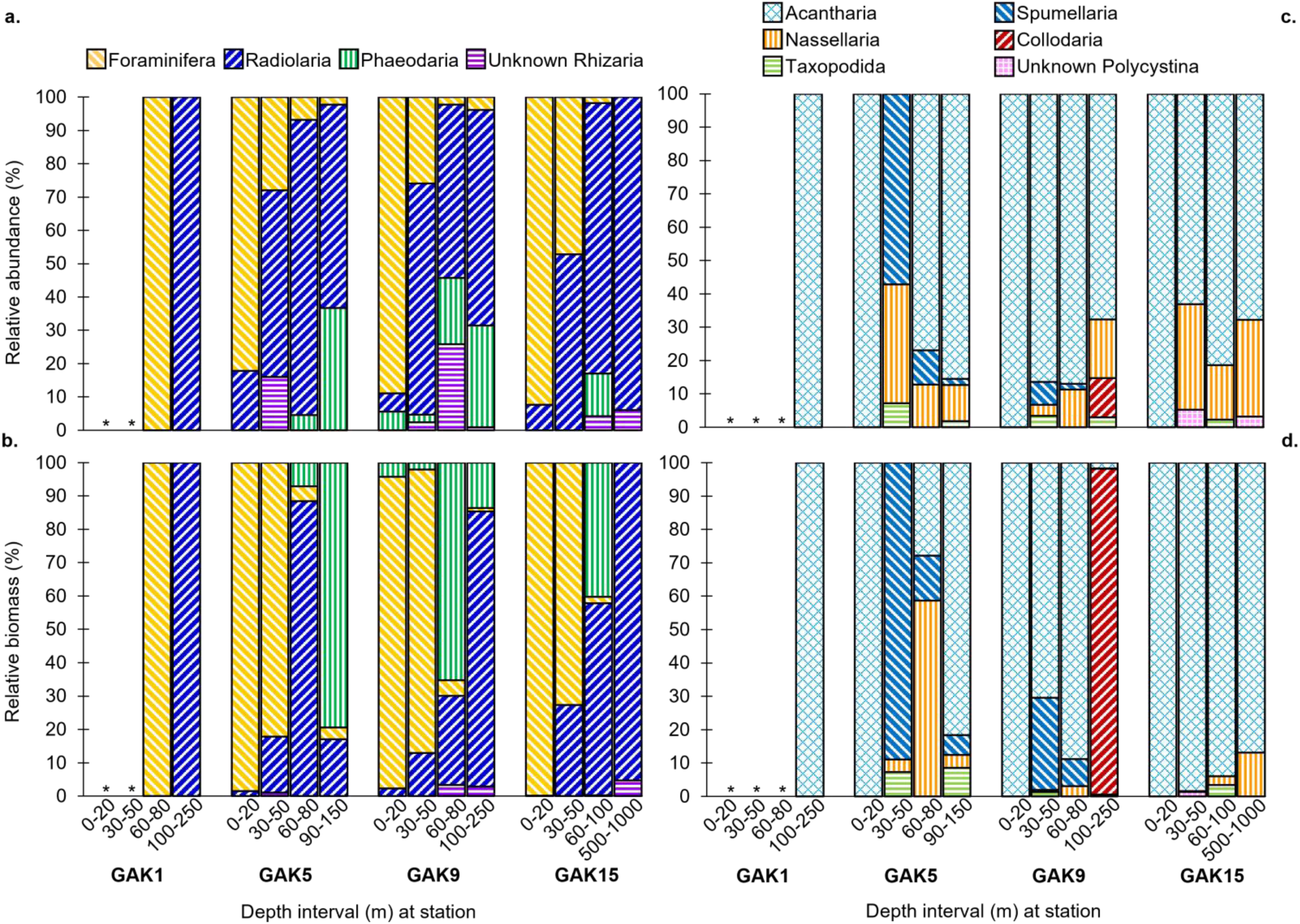
Percent composition of Rhizaria. a) abundance (cells L^-1^) (n=897) and b) biomass (ng C L^-1^) (n=818) by major taxonomic group and Radiolaria c) abundance (n=549) and d) biomass (n=507) by taxon at different depths across the Seward line. * = none detected.

NGA Rhizaria communities were more clearly separated by depth than by station as demonstrated by nMDS analysis. Surface communities were distinct from many of the deeper communities but similar among stations, as seen by their clustering at the extreme left of NMDS1 (Figure 11). In contrast, mid, intermediate, and deep communities were not clearly separated. In general, Rhizaria communities were not obviously separated by station as shown by their non-unique locations in ordination space.

**Figure 11.**
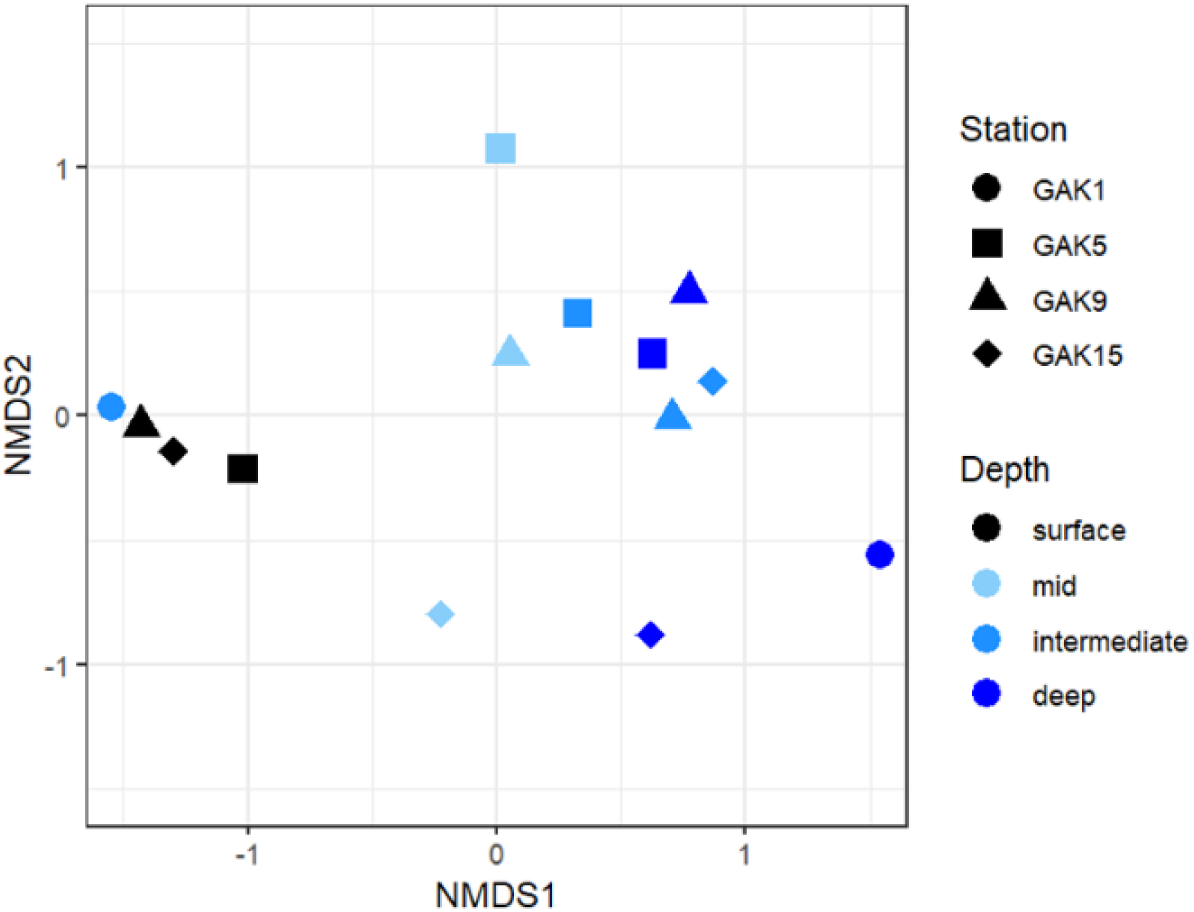
Relationship between Rhizaria communities at different station depths in ordination space.

### Trophic interactions

The proportion of Rhizarians with captured prey generally increased with distance offshore and declined with depth (Figure 12a). The highest proportions of Rhizaria with captured prey were observed far offshore at GAK15. Notably, incidences of prey capture were not observed nearshore (GAK1). Highest proportions of Rhizarians with captured prey were at or near the at the surface, with station maxima ranging from 25 to 33% of individuals (Figure 12a). The percentage of Rhizarians with prey captured reached a depth minimum of 3% in the deep waters at GAK15 but was also low in the intermediate and deep waters at GAK5 and GAK9 as well as at mid depths at GAK5 (Figure 12a).

**Figure 12.**
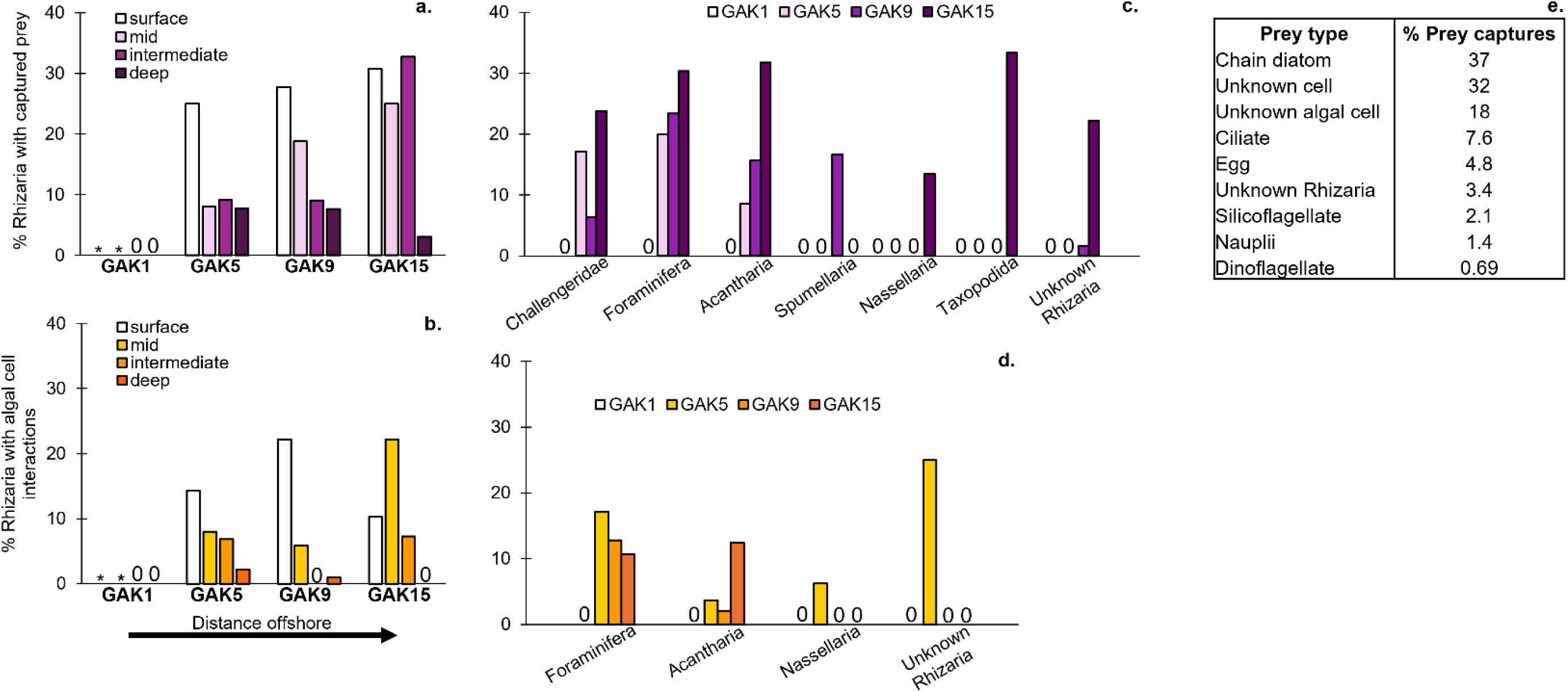
Summary of Rhizaria trophic interactions in the NGA. Distributions of % Rhizaria with **a)** captured prey and **b)** algal cell interactions at different depth intervals (see Supp. Table 1) across the Seward line (n=897). Grouped by taxon, distributions of % Rhizaria with **c)** captured prey and **d)** algal cell interactions across the Seward line (n=897). Taxa that lacked such interactions were not included. Total # of individuals with captured prey = 145 and with algae = 45. * = no Rhizaria detected; 0 = no individuals with captured prey (a and c) or algae (b and d). **e)** Table of prey types and the proportion each was involved in a prey capture event (n=155).

Most taxonomic groups identified in this study had individuals that exhibited prey capture. The proportion of individuals from a particular taxon with captured prey varied between 1.7% and 33% across the Seward line, although the majority of incidences at each station were >13% (Figure 12c). Highest prey capture incidence was seen at GAK15 for nearly all taxa (exception: Spumellaria), with Foraminifera, Acantharia, and Taxopodida reaching the highest levels (∼30%; Figure 12c). Foraminifera, Acantharia, and Challengeridae also captured significant prey at shelf stations GAK5 and GAK9. Diatoms (*Chaetoceros* spp., *Corethron* spp., *Melosirales* spp., and *Thalassiosira* spp.) were the most common prey item followed by cells of unknown identity (37 and 32% of prey captures, respectively, Figure 12e). Other types of prey (e.g. Figures 4b,c,f,g,m, 6z, 7b,c and 8d) were unidentified red and yellow-orange fluorescent algal cells, ciliates including tintinnids, copepod eggs, unknown Rhizarians, silicoflagellates, nauplii, and dinoflagellates (*Protoperidinium* spp.).

The proportion of Rhizarians that interacted with algae occurred at all stations except on the inner shelf at GAK1, and decreased with depth (Figure 12b). The highest incidences offshore were at the surface and mid depths (GAK5 to GAK15, range 14 to 22%, Figure 12b).

Foraminifera, Acantharia, Nassellaria, and Unknown Rhizaria exhibited interactions with algal cells. Epi-fluorescence images are shown in Figures 3f, 4d,h and 7a,d,e. Foraminifera had the highest proportions of individuals with algal cell interactions (GAK5 to GAK15, range 11 to 17%, Figure 12d) followed by Acantharia (GAK5 to GAK15, range 2.0 to 12%, Figure 12d). Incidences of algal cell interactions in Foraminifera and Acantharia demonstrated opposite cross-shelf trends. Proportions of Foraminifera with algal cell interactions decreased with distance offshore while proportions of Acantharia that interacted with algae generally increased with distance offshore. Nassellarians and Uknown Rhizarians with algal cells were only present at GAK5.

## DISCUSSION

Our study presents the first quantitative depiction of living planktic Rhizaria ecology in the eastern subarctic Pacific Ocean. Our Rhizaria abundance estimates are some of the highest reported from any ocean environment, peaking at 25 cells L^-1^. Therefore, we suggest a revision to the current biogeographical paradigm, which posits that abundances are highest at the equator and decrease towards the poles (Casey 1966, 1971, Petrushevskaya 1971, Renz 1976, Bé 1977, Boltovskoy and Correa 2017). In the NGA, Rhizaria were more likely to be found farther from shore. The major taxonomic groups exhibited clear vertical depth niches: Foraminifera preferred to live near the surface, Radiolaria were widespread, and Phaeodaria favored deep waters. Rhizaria occupy virtually all depths across the world’s ocean from the surface to the depths of ocean trenches. The longstanding view of Polycystine Radiolaria water column distributions is that abundances are highest in the upper 100 m and decrease with depth (Boltovskoy et al. 2017). In contrast, Foraminifera are reported to follow chlorophyll distributions and stay above the thermocline (Kimoto 2015), while Phaeodaria Cercozoans are the deepest ocean dwellers, usually residing below 300 m (Boltovskoy et al. 2017). In our study, Acantharia was a morphologically diverse, cosmopolitan group that dominated the Rhizaria community. An abundant new species was discovered that was deemed an Unknown Rhizarian, but we welcome other interpretations from fellow researchers. Our data supports the current understanding that Rhizaria are predators of common meso- and microplankton such as diatoms and ciliates and hosts to symbiotic/commensal algal cells, improving our understanding of the unique role that Rhizaria play in the food web dynamics of this productive marine region.

### Comparison of NGA Rhizaria ecology to other oceanic systems

This study is one of seven reported to date that used Niskin bottle samples to obtain intact Rhizaria (Michaels 1988, 1991, Gowing 1989, González 1992, Gowing and Garrison 1992, Michaels et al. 1995, Stoecker et al. 1996), and is the only from the North Pacific.

Plankton net mesh sizes used in other studies would have captured mainly large Rhizarians, missing smaller taxa, juveniles, and fragile taxonomic groups such as Acantharia. Imaging instruments also do not effectively identify small Rhizarians. The Niskin bottle - 50 µm mesh - reverse filtration method utilized here selected for the total Rhizaria size range; however, larger, rarer forms were not found given the relatively small volumes of water analyzed (∼9 L per sample). Here we primarily draw comparisons with studies that used Niskins, no matter the sampling location, but also those that employed net tows at comparable latitudes in the North Pacific.

### Northwest North Pacific Ocean

The NGA Rhizaria community was similar to that of the northwest North Pacific (NP) region in terms of composition and vertical depth partitioning but differed by abundances and species diversity. All three reports on Rhizaria planktonic distribution in the northwest NP, all of which used net tows, showed that Polycystine Radiolaria and Phaeodaria abundances generally increased with depth (Reshetnyak 1955, Okazaki et al. 2004, Ishitani and Takahashi 2007), whereas only a subset of Polycystines (Nassellaria) and Phaeodaria in the NGA followed this trend. In the Okhotsk Sea during late summer and along the Japan coast during late spring, abundances of these taxa were lower than in the NGA (Table 4).

**Table 4.** Comparison of Rhizaria abundances in different oceanic systems. Sampling method: N=Niskin, T=net tow, with analyzed volume (in parentheses) in liters, unless otherwise noted; nd=not determined.

| Location | Season | Method (volume) | Abundance (cells L<sup>-1</sup>) |  |  |  |  |  | Reference |
| --- | --- | --- | --- | --- | --- | --- | --- | --- | --- |
|  |  |  | Total Rhizaria | Acantharia | Foraminifera | Phaeodaria | Polycystina | Taxopodida |  |
| Okhotsk Sea | late summer | T (2.7 to 54 m<sup>3</sup>) | N/A | nd | nd | range 0 to <1 | range 0 to <1 | nd | Okazaki et al. 2004 |
| Japan coast | late spring | T (0.12 to 1.8 m<sup>3</sup>) | N/A | nd | nd | range 0 to 1.2 | <1 | nd | Ishitani and Takahashi 2007 |
| Weddell Sea | autumn | N (0.12 to 3.4)<br>T: <300 µm Phaeodaria (5 to 7);<br>>1mm Phaeodaria (166 to 600) | N/A | range <1 to 2.8 | <1 | <300 µm: range <1 to 3.1<br>>1 mm: <1 | range <1 to 2.2 | range <1 to 4.2 | Gowing 1989 |
| Weddell Sea | winter | N (60)<br>T: >1.6 mm Phaeodaria (100 m<sup>3</sup>) | N/A | range 0 to <1 | range 0 to <1 | <1.6 mm: range <1 to 2.7<br>>1.6 mm: range 0 to <1 | range 0 to <1 | range 0 to 2.8 | Gowing and Garrison 1992 |
| Equatorial Pacific | October | N (unknown) | N/A | avg. 8 | avg. 4.5 | nd | avg. 2 | nd | Stoecker et al. 1996 |
| Galapagos Vents | March | N (2 to 5) | N/A | max 29 | nd | nd | nd | nd | Michaels 1988 |
| N Pacific gyre, VERTEX station | multiple | N (10 to 12) | N/A | max 4.1 | nd | nd | nd | nd | Michaels 1991 |
| N Atlantic gyre, BATS site | spring, fall | N (12) | N/A | avg. 1.2 | avg. <1 | nd | nd | nd | Michaels et al. 1995 |
| N Gulf of Alaska | summer | N (9) | range 0 to 25 | range 0 to 12 | range 0 to 4.1 | range 0 to 3.1 | range 0 to 3.1 | range 0 to <1 | This study |

Rhizaria diversity was substantially greater in the northwest NP, on the order of dozens of species. In contrast, we identified 31 Polycystina morphotypes and only four Phaeodaria (three Challengeridae plus *P. acornis*) morphotypes. The greater diversity is almost certainly a consequence of using plankton nets, which filter large volumes of water and thus collect rarer taxa. These studies did not provide any data on Acantharia or Taxopodida Radiolarians for comparison. Most studies conducted in the North Pacific lack data pertaining to these major Radiolarian groups (e.g. Takahashi 1997, Okazaki et al. 2004, Itaki et al. 2008, Ikenoue et al. 2019, Bradshaw 1959, Petrushevskaya 1971, Beers et al. 1982, Renz 1976, Boltovskoy and Jankilevich 1985, Kling and Boltovskoy 1995).

### Southern Ocean

Rhizaria assemblages in the NGA and SO were similar in terms of species occurrence and depth distributions, but abundances and dominant species differed. Two studies that used a Niskin bottle sampling approach similar to ours quantified Rhizaria distributions in the Weddell Sea during austral autumn (Gowing 1989) and winter (Gowing and Garrison 1992). Total Rhizaria abundances during autumn were similar to the summer NGA, while winter abundances were 3x lower.

“Small” Phaeodaria (<300 µm in autumn sampled with net tows, and <1.6 mm in winter sampled with Niskin bottles) was one of the most abundant Rhizaria subgroups in the SO, a clear contrast with community structure in the NGA where Acantharia dominated. However, absolute abundances of small Phaeodaria were similar between the two high latitude oceans (Table 4).

Phaeodaria assemblages characteristically inhabit deep water with abundances generally increasing with depth in both the SO and NGA. The Challengeridae family comprised a major portion of Phaeodarians in both high latitude regions, and small Phaeodarians contributed similar biomass to the Rhizarian community in the SO winter (max 0.06 µg C L^-1^, Gowing and Garrison 1992) and NGA summer (max 0.02 µg C L^-1^, Challengeridae and *P. acornis*, Table 3). Even at depths where Phaeodaria made up fewer than 50% of the Rhizarian community by abundance in the NGA (Figure 10a), this group contributed substantially more biomass than abundant Acantharia Radiolarians because of their larger biovolumes. Most depths were influenced in this way by biomass of Challengeridae or *P. acornis* (Table 3).

Acantharia was substantially more abundant in the NGA compared to the SO. In the Weddell Sea, Acantharians were up to 4 to10x sparser during autumn and winter, respectively, than in the NGA (Table 4). Water column distributions were similar though; Acantharia generally increased with depth to ∼100 m in both regions, although their depth maximum was shallower (30-40 m) in the SO during winter. *S. zanclea* (Taxopodida) was another dominant species in the SO, a contrast with NGA abundances which were up to 4x less (Table 4). *S. zanclea* distributions increased with depth to ∼150-250 m similarly in both the Weddell Sea and NGA. Polycystina abundances and distributions did not vary much across these high latitude ecosystems.

Foraminifera abundances were consistently low in the Weddell Sea, while in the NGA this group was up to 4x greater (Table 4). Unlike the decline with depth seen in the NGA (Figure 9d), Foraminifera did not exhibit a clear vertical gradient in Antarctic waters.

In general, Antarctic Rhizaria abundances in the NGA summer and the SO autumn were similar. Although all major subgroups were present in each region, community composition differed. Most prominently, Acantharia and Foraminifera were more abundant in the NGA, while *S. zanclea* was more abundant in the SO; abundances of other taxa were similar between the two ecosystems. Because these marine systems share high latitude characteristics including moderate-to-high productivity summers, SO Rhizaria abundances might have exceeded those of the NGA if sampled during spring or summer.

### Low latitude oceans

NGA Acantharia and Foraminifera abundances were similar to those of the equatorial Pacific (EP; Stoecker et al. 1996), but Foraminifera biomass was greater in the NGA, which likely indicates prevalence of larger cells. Both regions shared Acantharia as the dominant taxon with similar biomass (upper 100 m, 79 ng C L^-1^, Table 3; upper 90 m, ∼110 ng C L^-1^, Stoecker et al. 1996), while NGA Foraminifera biomass was clearly greater (350 ng C L^-1^, Table 3; ∼80 ng C L^-1^, Stoecker et al. 1996). A similar size range of individuals was sampled in the NGA (>50 µm) and EP (>20 µm); most Foraminifera were >64 µm.

The depth-niche of Acantharia was deeper and more widespread in the NGA compared to another EP site, the Galapagos Vents (Michaels 1988), but abundances were lower in our study region (Table 4). Acantharia also dominated the Rhizaria community at the vent site.

Acantharia abundances in the upper 20 m were up to 30x higher (range 1.70-28.6 cells L^-1^) than in the NGA and declined with depth, in contrast to the NGA depth increase. Acantharia peak abundance at the vent site was 2x greater than in the NGA (Table 4). Despite major ecosystem contrasts, Acantharia dominance of the Rhizarian community was evident in the subpolar NGA and both EP sites.

There was a stark contrast in the abundances and habitat of Acantharians in the NGA versus the subtropical gyres of the central NP and North Atlantic (NA). In the NGA summer, maximum Acantharia abundance was deeper (∼60-100 m) and 3x higher compared to the NP gyre (Michaels 1991, Table 4). Lower abundances and a shallower depth niche for Acantharians were also seen in the NA central gyre during spring and fall (Michaels et al. 1995). NA central gyre peak Foraminifera abundances were 4x lower than in the NGA (Table 4) and no distinct depth niche was evident. Given these opposing vertical distributions and consistently lower gyre abundances, Acantharia and Foraminifera ecologies appear to differ between our mesotrophic subarctic study area and the oligotrophic subtropical gyres.

### Distribution patterns

NGA Rhizaria communities were structured according to depth during summer 2023. Planktic Foraminifera dominated the surface, Radiolaria exhibited a more cosmopolitan vertical distribution, and Phaeodaria usually resided below 60 m. Many researchers have attempted to connect Rhizaria vertical distributions to oceanographic parameters including temperature and salinity (Petrushevskaya 1971), carbon export (Guidi et al. 2016), potential prey (González 1992), salinity, chlorophyll-a, and dissolved oxygen (Ishitani and Takahashi 2007), and the Arctic Oscillation (Ikenoue et al. 2012). Others have suggested that distributions are influenced by seasonal fluctuations in the physical oceanography such as a water mass influx (Casey 1971, Björklund 1974). We hypothesize that the deeper-dwelling Radiolaria and Phaeodaria are most successful where they can avoid entanglement with abundant upper photic zone particles during the NGA’s productive summer season. In contrast, the surface habitat allows Foraminifera to maintain their interactions with algal cells. Radiolarians and Phaeodarians were mainly absent from depths above 30 m and 60 m, respectively. Whether this indicates they were retained in the same water mass across the gulf, or simply are not found in shallower waters because they do not thrive there, is unknown. Rhizaria abundances and biomass generally increased with distance offshore in the NGA. This supports the current theory that Rhizaria prefer open ocean salinities (Boltovskoy et al. 2017) and could explain their scarcity at low-salinity station GAK1.

Our data from the subarctic NGA contradicts the current understanding of Rhizaria biogeography. Rhizaria abundances and diversity are thought to be lowest at high latitudes and greatest in the equatorial regions (Casey 1966, 1971, Petrushevskaya 1971, Renz 1976, Bé 1977). However, we provide evidence to the contrary. Overall Rhizaria abundances in the NGA were similar to those at the equator. Moving northward to the subtropics, NGA abundances were markedly greater than those in the North Pacific and North Atlantic central gyres.

Furthermore, Rhizaria abundances were greater in the NGA than other ecosystems at comparable high latitudes, including the Southern Ocean (albeit at a different season) and northwest North Pacific Ocean. We propose a restructuring and refining of the Rhizaria equator-to-pole biogeographical gradient in the Pacific and Southern Oceans, as follows: abundances are high at the equator, decline in the subtropical gyres, and rise again with increasing proximity to the poles.

### Trophic activities

#### Prey interactions

Prey types and capture incidence for NGA Rhizaria resembled those reported by two studies conducted in other oceans. Acantharians collected from the Gulf Stream and Sargasso Sea primarily preyed on diatoms and ciliates including tintinnids, while Foraminiferans in these areas captured a wider range of prey items including diatoms, eggs, gelatinous/soft zooplankton, fecal pellets, ciliates including tintinnids, and other Rhizarians (Swanberg and Caron 1991). That study reported roughly 40-50% of individuals with prey captured, whereas we found lower capture frequencies (3-33%, Figure 12a) depending on sample location. SO Foraminifera, *S. zanclea*, and Phaeodaria held a variety of food items in digestive vacuoles such as diatoms, dinoflagellates, trichocysts and heterotrophic protists, <3 µm algae, Chlorella-like cells, bacteria, and other cellular or silicious fragments (Gowing 1986, 1989, Gowing and Garrison 1992). The same prey plus a few additional types were captured by NGA Rhizarians . We hypothesize that prey items like phyto- and microzooplankton remain consistent even from opposite sides of the world because they share a size range with Rhizaria, and their prevalence allows for high likelihood of capture.

Prey capture incidence in the NGA generally declined with depth and, even more strongly, increased with distance offshore. Prey capture was likely predominant at the surface because higher prey concentrations there would increase encounter rates with these passive suspension feeders. Maxima in >20 µm chlorophyll-a concentrations (representative of diatoms, a common prey type, as well as large dinoflagellates) were between 10-30 m on the shelf and at 40 m off the shelf (Figure 2f). These depths largely coincide with the 0-20 m depth interval that revealed the most individuals with prey captured. Because Rhizaria abundances were highest in deeper waters but prey capture incidence there was low, these organisms may rely on nutrition from detritus and marine snow particles to survive below the photic zone. High prey capture incidence coincided with low total chlorophyll at GAK15. Moreover, as was the case for all stations sampled, most of the chlorophyll originated from <20 µm algal cells which do not include common Rhizaria prey. Therefore, chlorophyll concentration was not a clear indicator of Rhizaria predation success, and it remains unclear why prey capture incidence was consistently high at GAK15 whilst prey availability there was low.

Challengeridae, Foraminifera, and Acantharia consistently displayed the highest incidences of prey capture. Phaeodaria are thought to be non-selective feeders or even detritivores, as they are often found with unknown organisms captured or digesting amorphous material (Anderson 1983, Gowing 1986). González (1992) observed that Phaeodaria were important minipellet producers in the SO. We have evidence to support this idea because 6-24% of Challengeridae (including *P. acornis*) had captured unknown algal material that fluoresced red to yellow, indicative of photopigments, inside the phaeodium (Figure 12c). There are few data on the trophic biology and predator-prey interactions of the above groups, so further research is necessary to understand if different Rhizaria subgroups are selective feeders, generalists, and/or detritivores, and whether they compete for the same prey items.

#### Algal cell interactions

The proportion of Rhizaria with associated algal cells was highest in shallow waters (<50 m) and increased with distance offshore, remaining under 30% overall (Figure 12b). Rhizarians that maintain algae must reside where there is sufficient light for photosynthesis. The cross-shelf pattern is a bit more difficult to decipher. The offshore NGA waters are typically characterized as a transition region between the shelf and fully iron-limited, HNLC waters of the subarctic gyre. Therefore, HNLC conditions can be detected near the seaward periphery of the NGA sampling region at GAK15 some seasons, as was the case during summer 2023.

Mixotrophic Rhizaria are important hubs of primary productivity in oligotrophic regions like the Sargasso Sea (Caron et al. 1995, Stoecker et al. 2017). Mixotrophy could be a favorable trait in the iron-limited offshore waters of the NGA, allowing Rhizaria to transition between nutrient acquisition modes depending on environmental conditions or prey availability. Microalgae in these resource-limited regions also benefit if the Rhizarian host provides a stable microenvironment with concentrated, nutrient-rich waste products. In general, lower productivity habitats might favor Rhizaria-algal associates given these mutual benefits.

Foraminifera and Acantharia were the only taxonomic groups consistently associated with algal cells in the NGA (Figure 12d); Acantharia had fewer incidences of algal interactions than Foraminiferans. Numerous Acantharia, Polycystina, and Foraminifera species harbor algal symbionts, whereas Phaeodaria are not known to partake in this trophic activity (Stoecker et al. 2009). Associated algal cells could be true endosymbionts, commensal or parasitic hitchhikers (Hemleben et al. 1989), prey, or even sequestered in an “inverted parasitism” relationship whereby only the host benefits (Decelle 2013). Takagi et al. (2019) identified species-specific differences in Foraminifera photosymbiosis and proposed, as did Stoecker et al. (2009), that there is a mixotrophy spectrum from heterotrophy to phototrophy with varying degrees of algal acquisition permanence, which is possibly dependent on host genotype, habitat, and/or symbiont physiology. Foraminifera algal associates include dinoflagellates as well as pelagophytes, chrysophytes, and *Synechococcus* spp. (Takagi et al. 2019), while Acantharia symbionts are usually prymnesiophytes (Stoecker et al. 2009). Unfortunately, we could not determine the algal taxa associated with NGA Rhizarians.

## Conclusion

This study provides a comprehensive view on Rhizaria ecology in the eastern subarctic North Pacific. Our new subpolar dataset challenges the current consensus on biogeographical distribution in the Pacific and Southern Oceans. The existing idea is that abundances are highest near the equator and decrease toward the poles. Our data support a restructuring of this gradient, whereby abundances are high at the equator, low in the subtropical gyres, and rise again with increasing proximity to the poles. This is an ambitious hypothesis due to the severe lack of research from different areas of the Pacific and Southern Oceans; for the data sets that do exist, contrasting methodologies and sampling seasons complicate direct comparisons. Many studies, including ours, lack replication and are temporally limited. Thus, there is much work to be done across all oceans to better comprehend the foundational aspects of Rhizaria biology and ecology.

## ACKNOWLEDGEMENTS

We thank Kaleigh Ballantine, Asher Marvy, and Katey Williams for sampling assistance and the crew of R/V *Kilo Moana* for making at-sea sample collection possible. This work was funded by the National Science Foundation (LTER grant 1656070), the North Pacific Research Board (grant 2108), and Western Washington University’s Graduate Research and Creative Opportunities Grant, Mickey and Carole Ghio Science Scholarship, and Marine and Coastal Science Program Graduate Student Support.

## SUPPLEMENTAL MATERIAL

**Supplemental Table 1.** Locations and depths for Rhizaria sampling (n=16).

| Station | Depth interval (m) | Depths sampled |
| --- | --- | --- |
| GAK1 | 0-20 | 0, 10, 20 |
|  | 30-50 | 30, 40, 50 |
|  | 60-80 | 60, 70, 80 |
|  | 100-250 | 100, 150, 250 |
| GAK5 | 0-20 | 0, 10, 20 |
|  | 30-50 | 30, 40, 50 |
|  | 60-80 | 60, 70, 80 |
|  | 90-150 | 90, 100, 150 |
| GAK9 | 0-20 | 0, 10, 20 |
|  | 30-50 | 30, 40, 50 |
|  | 60-80 | 60, 70, 80 |
|  | 100-250 | 100, 150, 250 |
| GAK15 | 0-20 | 0, 10, 20 |
|  | 30-50 | 30, 40, 50 |
|  | 60-100 | 60, 80, 100 |
|  | 500-1000 | 500, 750, 1000 |

**Supplemental Table 2.**
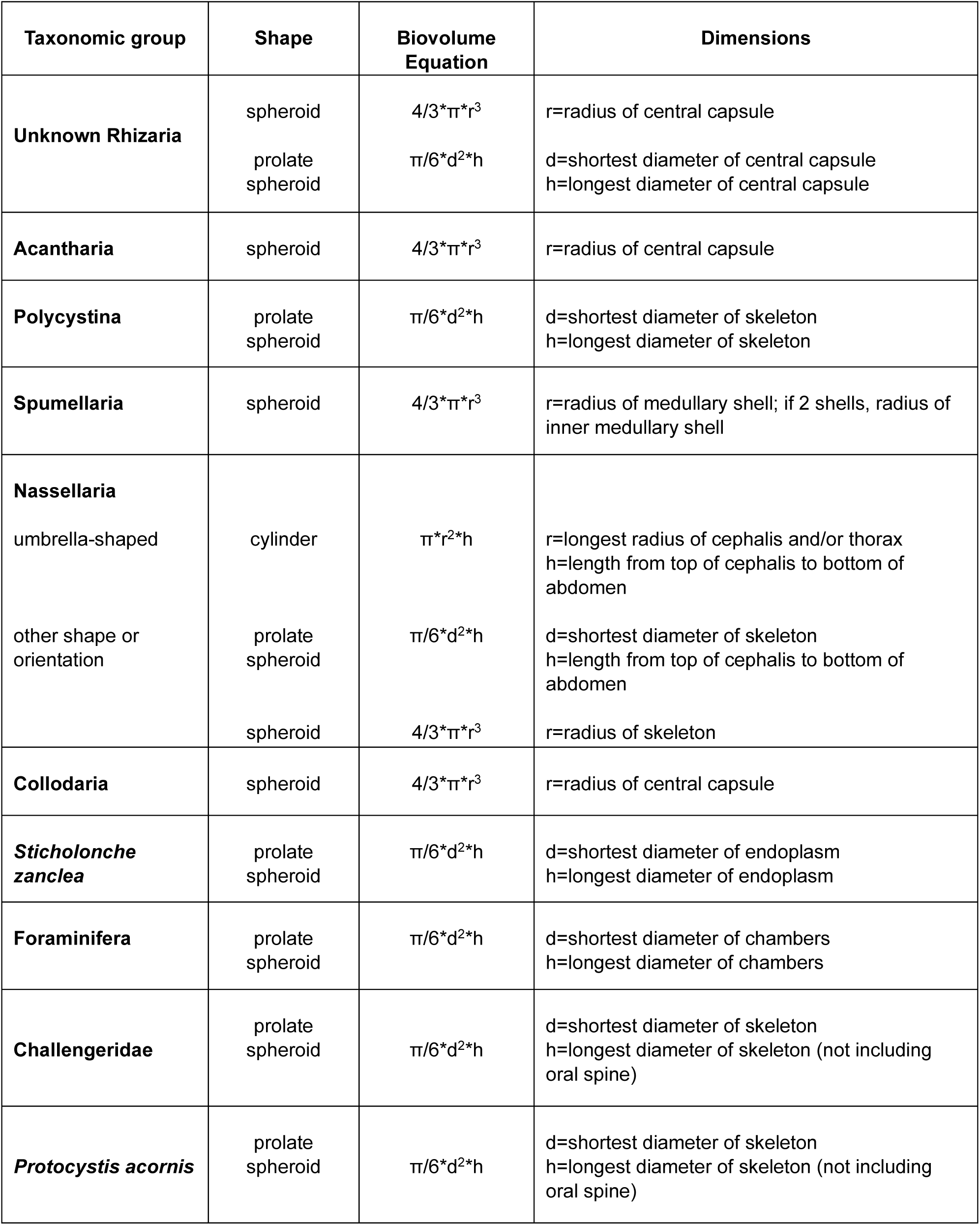
Biovolume measurement details.

## Notes

https://doi.org/10.6073/pasta/74f978401b5a224200a54ee5bf800165

### Competing Interest Statement

The authors have declared no competing interest.

